# Chromatin-sensitive cryptic promoters encode alternative protein isoforms in yeast

**DOI:** 10.1101/403543

**Authors:** Wu Wei, Bianca P. Hennig, Jingwen Wang, Yujie Zhang, Ilaria Piazza, Yerma Pareja Sanchez, Christophe D. Chabbert, Sophie H. Adjalley, Lars M. Steinmetz, Vicent Pelechano

**Author notes:** These authors contributed equally to the work (Wu Wei, Vicent Pelechano).

## Abstract

Cryptic transcription is widespread and generates a heterogeneous group of RNA molecules of unknown function. To improve our understanding of cryptic transcription, we investigated their transcription start site usage, chromatin organization and post-transcriptional consequences in *Saccharomyces cerevisiae*. We show that transcription start sites (TSSs) of chromatin-sensitive internal cryptic transcripts retain comparable features of canonical TSSs in terms of DNA sequence, directionality and chromatin accessibility. We define the 5’ and 3’ boundaries of cryptic transcripts and show that, contrary to RNA degradation-sensitive ones, they often overlap with the end of the gene thereby using the canonical polyadenylation site and associate to polyribosomes. We show that chromatin-sensitive cryptic transcripts can be recognized by ribosomes and may produce truncated polypeptides from downstream, in-frame start codons. Finally, we confirm the presence of the predicted polypeptides by reanalyzing N-terminal proteomic datasets. Our work suggests that a fraction of chromatin-sensitive internal cryptic promoters are in fact alternative truncated mRNA isoforms. The expression of these chromatin-sensitive isoforms is conserved from yeast to human expanding the functional consequences of cryptic transcription and proteome complexity.

## Introduction

Genomes are pervasively transcribed, producing a wide diversity of coding and non-coding RNAs (reviewed in (Jensen et al. 2013; Wei et al. 2011; Pelechano and Steinmetz 2013; Kaikkonen and Adelman 2018)), raising the question of the biological significance of such transcriptional activity (Jensen et al. 2013). Some of those transcripts are functionally relevant, such as the well-characterized long non-coding RNAs, antisense transcripts or alternative isoforms (reviewed in (Jensen et al. 2013; Pelechano and Steinmetz 2013; Pelechano 2017; Kaikkonen and Adelman 2018)). However it remains unclear which fraction of these transcripts exert a biological role (direct or regulatory). This question is particularly difficult to address when these transcriptional units arise within, or in close proximity to protein coding genes in the same strand. Thus their transcription signals are difficult to distinguish from the nearby or even overlapped protein coding genes. Among pervasively produced transcripts, so-called cryptic transcripts constitute a particularly heterogeneous group. Cryptic transcription is typically defined as the production of non-canonical transcripts of unknown function (Wei et al. 2011), where canonical transcripts can be interpreted as those encoding a full-length functional protein. The breadth of this definition shows that, despite their abundance and potential relevance for gene expression, our knowledge of this process remains limited.

Cryptic transcripts can be classified according to the mechanisms by how cells control their abundance: Cryptic transcripts levels may be modulated either by restricting transcription initiation or by selectively degrading them (reviewed in (Jensen et al. 2013)). For simplicity, we will refer to the first class of processes as “chromatin-sensitive” and to the second class as “RNA degradation-sensitive”. A classical example of chromatin-sensitive mechanisms is the emergence of cryptic transcripts from within gene bodies when histone deacetylation patterns are disrupted. Specifically, interfering with the activity of the Rpd3S deacetylase complex, which recognizes Histone 3 Lys36 trimethylation (H3K36me3) deposited by the histone methyltransferase Set2 during RNA polymerase II elongation, leads to intragenic cryptic transcription (Carrozza et al. 2005; Lickwar et al. 2009; Churchman and Weissman 2012; Chabbert et al. 2015; Malabat et al. 2015; Kim et al. 2016). Likewise, modulating nucleosome positioning by impairing the function of histone chaperons such as Spt6p also leads to the appearance of intragenic cryptic transcripts (Kaplan et al. 2003; Doris et al. 2018). Spt6p depletion causes decreased expression of most genic promoters while increasing the expression of intragenic ones, thus suggesting a potential competition for initiation factors (Doris et al. 2018). In contrast, the second class of cryptic transcripts (RNA degradation-sensitive) are constitutively produced and degraded by the cell, and thus become detectable only when RNA degradation is impaired (Jensen et al. 2013). For instance, cryptic unstable transcripts (CUTs) are identified in mutant cells with depletion of the nuclear RNA exosome (e.g. *rrp6∆*) (Xu et al. 2009; Neil et al. 2009).

Due to their proximity to or even overlap with protein-coding genes, dissecting the function of cryptic transcription units is especially complicated. In some contexts, cryptic transcription has been as sociated to “opportunistic transcription”, whereby RNA polymerase II is recruited to any open chromatin region, generating spurious molecules. However, annotating cryptic transcripts as functional or spurious is not trivial. This has been exemplified in multiple instances where either the RNA product itself or the transcriptional activity per se may have a clear functional impact. For example, the act of transcription itself can regulate the expression of neighboring genes through chromatin modulation (Martens et al. 2004; Hainer et al. 2011; van Werven et al. 2012; Kim et al. 2012; Xu et al. 2011; Chia et al. 2017; Brown et al. 2018). On the other hand, previous reports have shown that cryptic promoters can drive the expression of alternative isoforms with different post-transcriptional regulation or even encode alternative protein isoforms (Cheung et al. 2008; Arribere and Gilbert 2013; Pelechano et al. 2013; Fournier et al. 2012; Lycette et al. 2016; Carlson et al. 1983; Gupta et al. 2014).

To improve the classification of such events and further improve our understanding of cryptic transcription, we performed a comprehensive characterization of cryptic promoters in *Saccharomyces cerevisiae*. We carried out the analysis of both the biogenesis of cryptic transcripts and their post-transcriptional life with a focus on those derived from chromatin-sensitive mechanisms (*i.e. set2∆, rco1∆* and *eaf3∆*). As a comparison, we also examined the biogenesis of the RNA degradation-sensitive CUTs (*rrp6∆*, RNA degradation sensitive). We identified their transcription start sites (TSSs), and investigated their sequence preference and chromatin organization. To assess the post-transcriptional life of cryptic transcripts and better define their boundaries, we examined the association between TSS and polyadenylation site usage by TIF-seq (Pelechano et al. 2013). To investigate their coding potential, we performe d polyribosome fractionation followed by 5’ cap sequencing to investigate the association of cryptic transcripts with polyribosomes. We examined the ribosome protection pattern of cryptic transcripts measured by 5PSeq (Pelechano et al. 2015) focusing on the signature associated to internal methionine codons predicted to act as novel start codons. Finally, we validate our prediction using available N-terminal Mass Spectrometry data (Varland et al. 2018). Our work aims to investigate the functional relevance of chromatin-sensitive cryptic transcripts.

## Results

### Chromatin-sensitive and RNA degradation-sensitive cryptic transcripts show distinct TSS profiles

To understand how cryptic transcripts are generated, we performed a genome-wide mapping of their transcription start sites (TSSs) in *Saccharomyces cerevisiae*. We conducted 5’cap sequencing (Pelechano et al. 2016), which enables a precise identification of the 5’ end of transcripts in a wild-type strain (BY4741) and multiple mutants associated with cryptic transcription (Fig. 1A). To illustrate chromatin-sensitive cryptic transcription, we examined the TSSs profile of cells lacking Set2, the histone methyltransferase responsible for the co-transcriptional deposition of H3K36me3 (Carrozza et al. 2005). We also investigated the TSS profile of strains deficient in Rco1 and Eaf3, components of the Rpd3S histone deacetylase complex acting downstream of Set2. Furthermore we examined the emergence of cryptic TSSs in cells deficient for Set1, the histone methyltransferase responsible for H3K4 methylation and associated with cryptic transcription from promoter-proximal regions (van Werven et al. 2012; Kim et al. 2012). Finally, we conducted a comparative analysis of the TSS profiles of cryptic unstable transcripts (CUTs) that emerge upon depletion of the nuclear RNA exosome subunit Rrp6 (*rrp6∆*) (Xu et al. 2009; Neil et al. 2009) as an example of RNA degradation-sensitive cryptic transcription.

**Figure 1.**
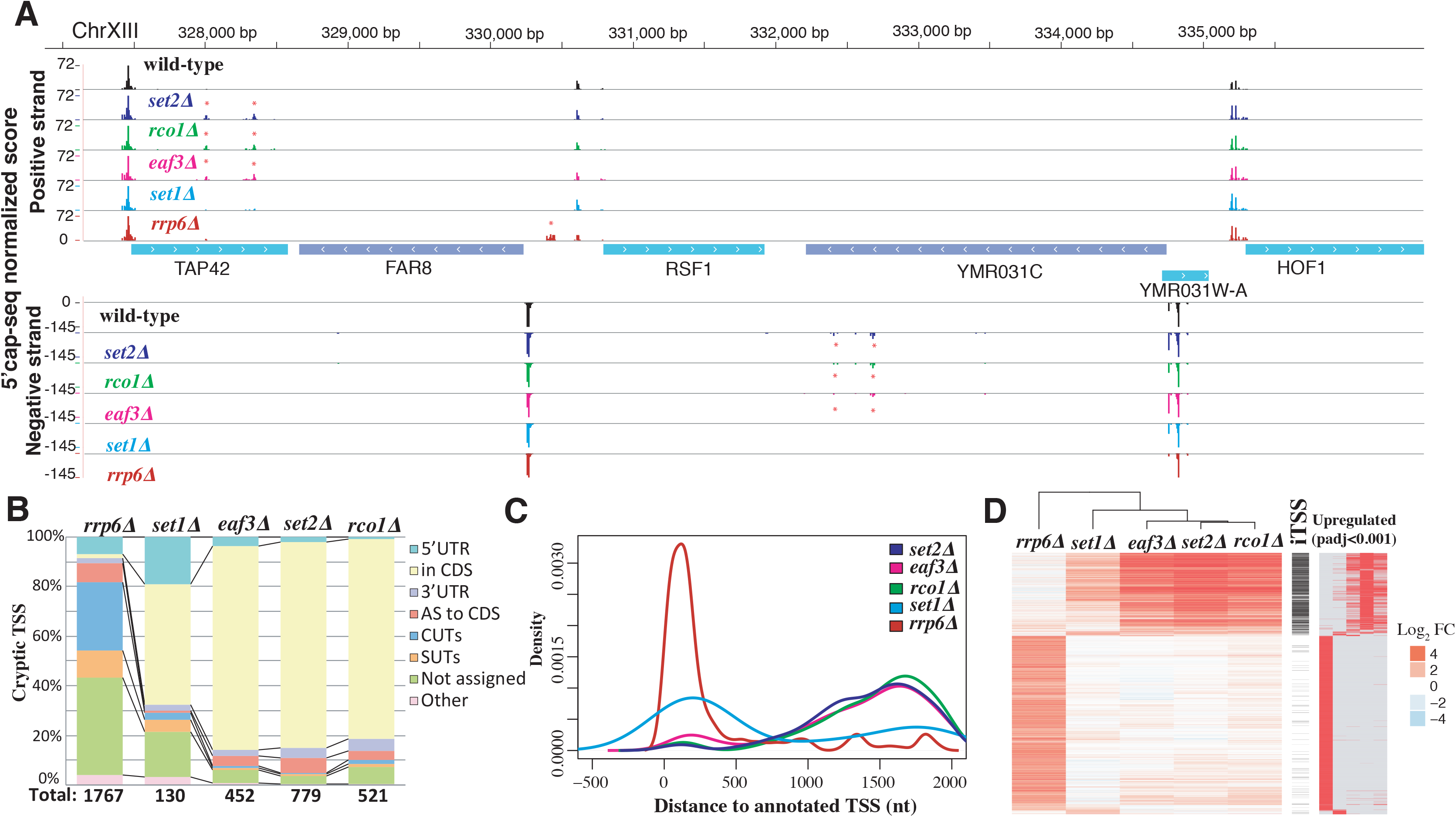
Genome-wide identification of chromatin and RNA-degradation sensitive TSSs. Detected chromatin-sensitive cryptic transcripts tend to overlap coding genes in the same orientation. (A) Representative 5’cap sequencing track. Score (normalized counts) of collapsed replicates is shown (see methods). Significantly differential expressed TSSs clusters marked by * (p-adj <0.001). (B) Classification of differentially expressed TSSs in respect to annotated features. Annotation of SUTs (Stable Unannotated Transcripts), CUTS and UTR lengths are from (Xu et al. 2009) (C) Distribution of differentially expressed TSSs in respect to annotated ORF-T TSSs. ORF-T refer to transcripts associated to canonical ORFs as described by strand-specific tiling arrays in (Xu et al. 2009) (D) Relationship between TSSs identified in the analyzed strains. Each horizontal line represents an identified TSS cluster. On the left side we display the relative fold change enrichment (FC) respect to the wild-strain in Log_2_ (red (up regulated) to blue (down regulated)). In black we indicate which of those identified TSS can be classified as iTSS. Finally, significantly differentially expressed TSSs comparing to wild-type are shown at the right (in red). Only TSSs identified as differentially expressed with respect to the wild type in at least one condition are shown.

In total, 44963 TSS clusters were identified across all datasets (Supplemental table S1). We used information from our biological replicates and unique molecular identifiers (UMIs) to identify differentially expressed TSSs across strains (adjusted p-val<0.001; see methods) (Fig. 1B and S1A). The disruption of the nuclear exosome (*rrp6∆*) led to the highest number of up-regulated TSS clusters in comparison to wild-type (1767), while deletion of *SET1* had a moderate effect (130 TSS up-regulated clusters). The other mutant strains, *set2∆, rco1∆* and *eaf3∆*, presented an intermediate phenotype (*i.e.* with 779, 521 and 452 up-regulated TSS clusters, respectively). Up-regulated *rrp6∆-sensitive* TSSs were detected in close proximity to the annotated TSSs of coding genes (often in opposite orientation to annotated TSSs (Xu et al. 2009; Neil et al. 2009)) while *set2∆-, rco1∆-* and *eaf3∆*-sensitive TSSs occurred preferentially within the body of genes (Fig. 1C and S1B) (Carrozza et al. 2005; Lickwar et al. 2009). Strains with mutations affecting the same pathway (*e.g. set2∆, rco1∆* and *eaf3∆*) shared a high number of up-regulated cryptic TSSs, while cryptic TSSs resulting from disruption of the nuclear exosome (CUTS, *rrp6∆*) occurred mainly outside of the coding regions (Fig. 1D and S1C). We then characterized the intragenic upregulated TSSs (iTSSs) that occur inside the coding region of genes and mostly originate from the Set2-Rco1-Eaf3 pathway (Fig. 1D). Our strand-specific detection approach enabled us to determine that most chromatin-sensitive cryptic iTSS are expressed in the same orientation as the corresponding ORF. This contrasts with what is observed for the RNA degradation-sensitive ones that arise more often antisense to the CDS than in the same orientation (red vs yellow in Fig. 1B). Previous strand specific RNA-seq analysis of the *set2∆* strain has identified the presence of internal Set2-repressed antisense transcripts (SRATs) (Venkatesh et al. 2016). Our work confirms their finding (SRATs displayed in red in Fig. 1B, S1A, S1E and S2), but further reveals that the vast majority of stable cryptic transcription overlaps the main transcript in the same orientation (yellow in Fig. 1B), a feature difficult to detect with conventional RNA-seq. To investigate the origin of the directionality of the chromatin-sensitive cryptic iTSS, we reanalyzed NET-seq (Churchman and Weissman 2012), RNA-seq (Venkatesh et al. 2016) and alternative TSSs datasets (Malabat et al. 2015) in addition to our data (Fig. S1D-E and S2). This revealed that although nascent transcription arises bidirectionally from cryptic promoters, cryptic transcripts in the same orientation as the main ORF are more stable and thus accumulate to a higher level. In fact, chromatin-sensitive iTSSs can also be detected, albeit at a much lower level, in wild-type conditions (see below). The Winston lab has recently investigated the appearance of intragenic promoters upon Spt6p depletion (*spt6-1004*) (Doris et al. 2018). We compared up to what degree *spt6-1004* up regulated intragenic promoters overlap with the chromatin-sensitive cryptic iTSS defined in this study (Fig. S3). As can be observed, chromatin-sensitive cryptic iTSS are only slightly increased in *spt6-1004*, while the vast majority of *spt6-1004* up-regulated intragenic promoters are not up regulated in a *set2∆* strain (Fig. S3A). Additionally, *spt6-1004* has a clear effect decreasing the expression of canonical genic promoters, while *set2∆* has a more punctuated effect in the body of the genes (Fig. S3B). This suggest that, although related, this two pathways control different subsets of cryptic promoters that are only partially overlapping. To gain a better understanding of the regulation of the chromatin-sensitive iTSSs, we decided to focus our analysis on those iTSSs occurring in the same orientation as their overlapping coding gene.

### Characterization of cryptic iTSS promoters

After identification of the putative promoter regions with cryptic iTSSs, we compared these with the canonical TSSs of protein-coding genes. iTSSs in all analyzed strains present a similar sequence composition to canonical TSSs, with a pyrimidine enrichment at the −1 and adenine at the 0 and −8 position (Zhang and Dietrich 2005; Pelechano et al. 2013) (Fig. 2A and S4A). Please note that transcript position 0 as refered here (first nucleotide of the transcript) is traditionally referred also as +1, when using a scale without 0. It is also important to note that molecules derived from cryptic iTSSs can also be detected in wild-type cells, although at a lower level (Fig. S1D). This suggests that cryptic iTSSs are used by at least a fraction of cells in normal growing conditions.

**Figure 2.**
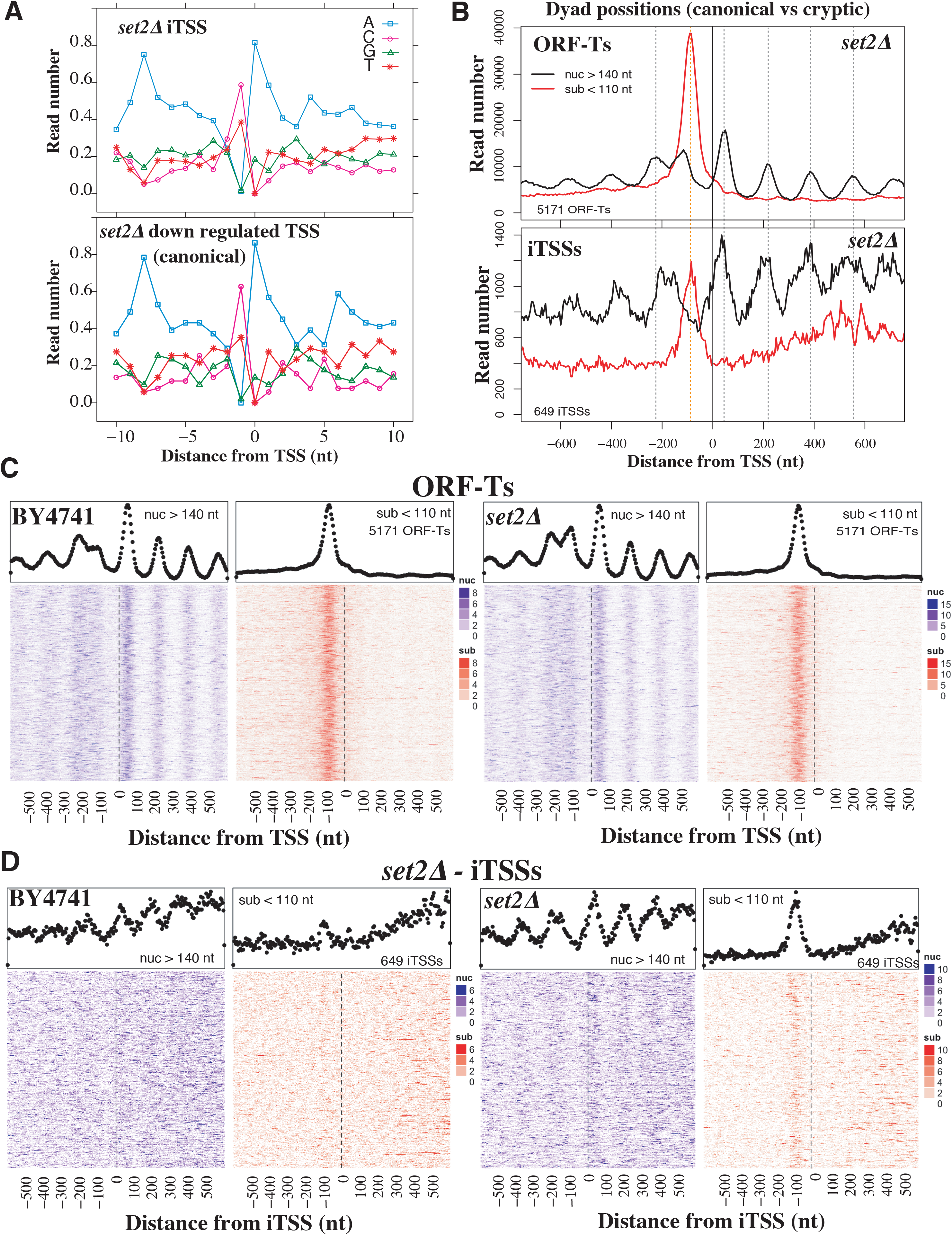
The sequence and chromatin features of iTSSs resemble those of canonical TSSs. (A) Sequence preference of *set2∆* iTSSs compared to canonical TSSs (*set2∆* down regulated that often overlap with canonical TSSs). (B) MNase protection pattern for canonical ORF-T TSSs. MNase fragments are distributed in nucleosome protection fragments (nuc) and sub-nucleosomal ones (sub) according to their length. Vertical dotted lines depict canonical dyad nucleosome axis (in black) and putative TF binding sites (in read). (C) Heatmaps depicting in detail the MNase protection pattern for canonical ORF-T TSSs in the wild-type strain and *set2∆*. Each line of the heatmaps correspond to an analysed region for nucleosome fragments (in blue) and subnucleosomal fragments (in red) ordered by gene expression (Xu et al. 2009). The metagene with aggregation of all the heatmap information is shown above in black dots. (D) Heatmaps depicting in detail the MNase protection pattern for set2∆ iTSSs as in C. Chromatin data is reanalysed from (Chabbert et al. 2015). Heatmap sorted by iTSS expression level.

Given that chromatin-sensitive iTSSs resemble canonical gene-coding TSSs in their base composition and directionality, we assessed whether this also applies to their chromatin organization. We used information on nucleosomal and subnucleosomal fractions from our previous high-throughput ChIP-seq experiments (Chabbert et al. 2015; 2018) (Fig. 2C, S4B and S5A) to analyse the MNase protection pattern around cryptic iTSSs. Cryptic iTSSs present the same MNase protection architecture as canonical TSSs, with an organized nucleosome array downstream of the TSS and a sub-nucleosomal protection site overlapping the region where TFs would typically associate (Henikoff et al. 2011)(Fig. 2D and S5B). This is particularly evident for the Set2-Eaf3-Rco1 sensitive iTSSs, as they are further away from canonical TSSs (Fig. 1C) and thus easier to disentangle from the MNase pattern associated to canonical promoters (Fig. 2D). A similar, although more discrete pattern (*i.e.* nucleosome array and upstream sub-nucleosomal protection pattern) can also be observed around the same iTSSs in the wild-type strain (Fig. 2D and S5B). The subnucleosomal fragments are only apparent when analysing whole cell extract, and are depleted after histone imunoprecipitation (Fig. S6A). This suggests that either histones are not bound to those fragments, or that they cannot be efficiently immunoprecipitated in our experimental conditions. The distance between the iTSS and the first nucleosome downstream (analogous to the +1 nucleosome) is similar to the distance present in canonical TSSs and the dyad axis (Fig. 2B and S5). However, the nucleosome-depleted regions (NDR) commonly associated to promoters are a bit smaller in the case of the “cryptic promoters” of iTSSs. In our experimental conditions we estimate that canonical NDR are approximately of 275nt, while iTSS NDR are 215nt and the distance between +1/+2 nucleosome dyads is of 165nt (Fig. S5). The presence of a periodic nucleosome organization in gene bodies around an internal “nucleosome depleted region” upstream of the cryptic iTSS, suggest that iTSSs tend to occur or contribute to synchronizing, regular nucleosome arrays that are detectable even in mixed cell populations. This, together with the detection of a basal level of cryptic iTSS expression (Fig. S1D), suggests that a small proportion of cells are expressing these cryptic transcripts even under normal conditions. It is important to note that iTSS NDR are longer than the average distance between nucleosome pairs even in a wild-type strain (Fig. S5B). This suggests that factors or genome features may actively make these internucleosome regions distinct. Additionally, our observation that cryptic iTSSs may be bound by TF at low levels even in normal conditions, is in agreement with recent evidence suggesting that TF such as Gcn4 can also bind and activate internal promoters (Rawal et al. 2018).

To confirm that cryptic iTSSs present the canonical marks associated with promoter activity, we analyzed other chromatin features. We focus on chromatin-sensitive iTSSs that are in general are distant from the canonical TSSs, and thus not obscured by canonical promoter marks (Fig. 1 C). We observed an increased signal of H3K4me3 at the first nucleosome (+1 nucleosome) downstream of the iTSSs in *set2∆* that decreases downstream of the cryptic promoters (Fig. S6B). As expected this is only apparent in this mutant strain as cryptic transcripts are expressed at a sufficient level to be detectable.

### Post-transcriptional life of iTSSs derived transcripts

Once confirmed that iTSSs present a canonical promoter structure, we sought to determine the complete length of the transcripts derived from iTSSs in order to gain information on their post-transcriptional life. We applied our previously developed Transcript Isoform Sequencing (TIF-seq (Pelechano et al. 2013)) approach that allows to jointly and unambiguously determine the start and end sites (TTSs) of each RNA molecule within a sample. We thus identified the start and end sites of all transcripts, including the chromatin-sensitive transcripts that initiating from iTSSs. We further compared the TSSs and TTSs of iTSSs initiated transcripts to those of canonical transcripts. We identified that most transcripts originating from an iTSS in the *set2∆* strain use the same polyadenylation sites as the canonical mRNAs. This was observed at both individual and genome-wide levels (Fig. 3).

**Figure 3.**
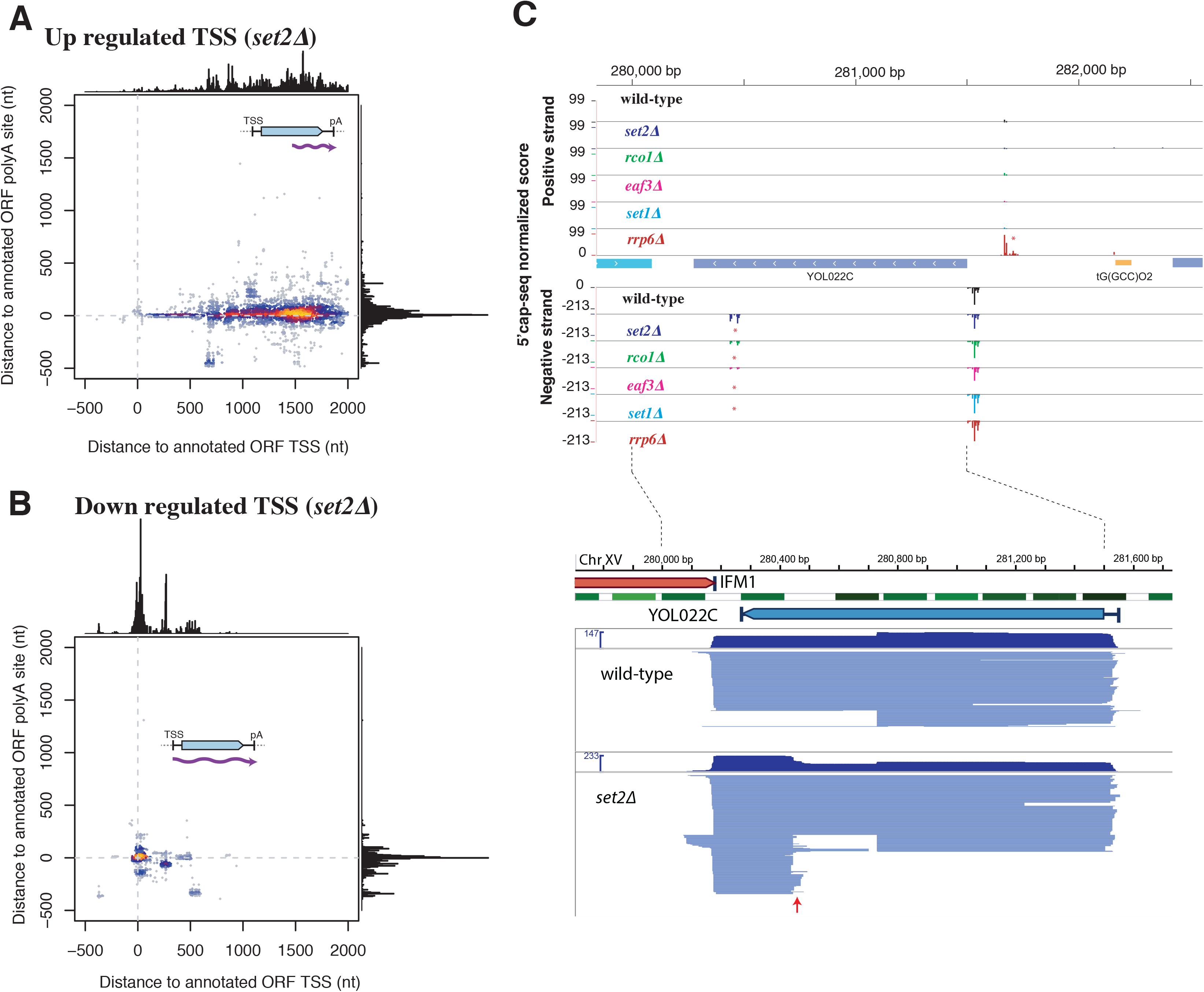
Full length of *set2∆* iTSSs derived transcripts use canonical polyadenylation sites. (A) The transcript start and end sites comparison between *set2∆* iTSSs initiated transcripts and annotated ORF-T boundaries (Xu et al. 2009). *set2∆* iTSSs derived transcripts originate within the body of the gene (internal 5’) but use canonical 3’ polyadenylations sites. (B) Down-regulated TSSs in *set2∆* use canonical 5’ and 3’ sites. (C) Example of TIFSeq coverage for YOL022C gene as an example. The upper part shows TSS mapping (as in Fig. 1A). In the bottom part we show full-length transcript in blue. Each line connecting between one identified TSS and poly(A) site represents one full-length transcript. The red arrow, indicated the appearance of a set2∆ sensitive iTSS. Nucleosomes are showed in green (Venters and Pugh 2009).

Specifically, most transcripts emerging from an iTSS in s*et2∆* originate within the gene body but use the canonical polyadenylation sites (Fig. 3A). This confirms and expands previous evidence from northern blot analysis (Kaplan et al. 2003; Carrozza et al. 2005). In contrast, TSSs down-regulated in *set2∆*, that in the vast majority correspond to canonical mRNA TSSs, generate transcripts that also use the canonical polyadenylation sites (Fig. 3B) (Xu et al. 2009). These suggest that stable chromatin-sensitive cryptic transcripts have the potential to encode N-terminal truncated proteins. As most chromatin-sensitive cryptic iTSSs can produce 5’ truncated mRNAs, we further investigated if they are associated with ribosomes. This is particularly interesting as those molecules are present at low levels even in wild-type conditions, which could function as alternative mRNA isoforms. Additionally, previous work showed that a fraction of internal cryptic transcripts is degraded thought Non-sense Mediated Decay (NMD), and thus putatively interact with the translation machinery enough to be surveyed by NMD (Malabat et al. 2015). To measure association with ribosomes of the stable chromatin-sensitive cryptic transcripts, we combined isolation of polyribosomes by sucrose fractionation with 5’ cap sequencing (Supplemental Table S2). As expected ORF-Ts TSSs are associated with polyribosome fractions, while non-coding RNAs such as SUTs (Stable Unannotated Transcripts) or CUTs are much less associated (Xu et al. 2009) (Fig. 4A and S7A-B). It is important to note, that although the bulk of CUTs and SUTs are not preferentially associated to ribosomes, a fraction of them could encode peptides (see below). mRNA molecules originating from chromatin-sensitive cryptic iTSSs are also enriched in the heavy polyribosome fractions that are associated with active translation. And this association does seem to depend on the length of the cryptic 5’UTR (Fig. S7C). This suggests that cryptic transcripts, especially those originating from chromatin-sensitive cryptic promoters, associate with ribosomes and have the potential to produce truncated proteins.

**Figure 4.**
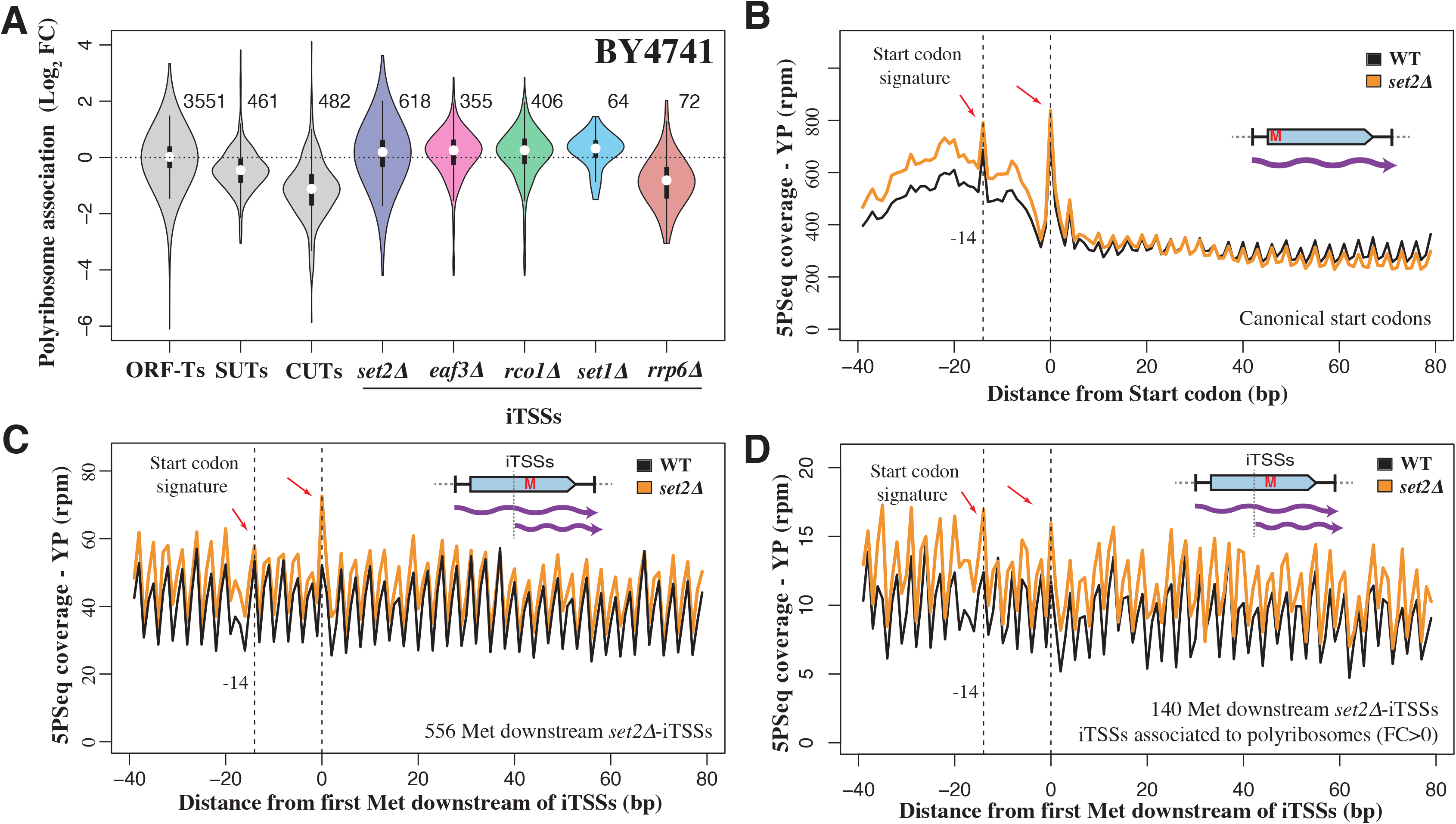
A fraction of iTSSs derived transcripts associate to ribosomes and the internal methionine can be recognized as a novel start codon. (A) Relative association with polyribosome fraction after sucrose fractionation versus total extract. Analyzed events (present at a sufficient level in the wild-type strain) are indicated to the right of each plot. (B) Example of 5PSeq start-codon associated signature after glucose depletion for coding genes. To decrease the effect of potential outliers, we assigned a value corresponding to the 95th percentile to values that were over this threshold at each distance from the start codon. (C) Start-codon associated signature after glucose depletion for predicted novel start codons in *set2∆* iTSSs derived transcripts. Those positions are expected to behave as internal methionines in a wild-type strain. (D) As in C, but showing the subset of cryptic start-codons which mRNAS as more associated to polyribosomes (fold-change >0 in A).

To assess whether ribosome-bound cryptic transcripts also are engaged in active translation, we assayed the ability of ribosomes to recognize such cryptic transcripts. To this aim, we used our previously developed 5PSeq approach, that measures ribosome dynamics by sequencing co-translational mRNA degradation intermediates (Pelechano et al. 2015; 2016). We have previously shown that yeast cells in slow growth conditions such as growth in minimal media or stationary phase present a characteristic ribosome protection pattern at the translation start codon consistent with inhibition of translation initiation (Pelechano et al. 2015; Pelechano and Alepuz 2017). To distinguish the translation of the canonical full length mRNAs from the shorter overlapping transcripts derived from iTSSs, we applied 5PSeq in glucose starvation to test if iTSSs initiated transcripts show translation start codon pattern (Zid and O’Shea 2014). In fact, we identify a 5PSeq protection pattern at −14 nt and at the start codon (Fig. 4B and S8). Initially, we tried to enhance the start codon signature using cycloheximide treatment, as it leads to a sharp increase of protection at −14 nt. However, as expected for an inhibitor of translation elongation, cycloheximide also leads to a massive increase of internal 5PSeq protection that obscure the signature of any internal cryptic translation start site (Fig. S8E-F). To enhance the observed start codon signature, we exposed cells to a glucose-free media for 5 minutes. By limiting translation initiation we increased the start codon signature and allowed the ribosomes engaged in translation *to run-off* the mRNA (Zhang Y and Pelechano V*, in preparation*), an effect that can be readily observed at the canonical start codons of annotated protein-coding genes (Fig. 4B and S8 A-B). We then analyzed the ribosome pattern associated with internal methionines and focused on those in-frame that could potentially be recognized as new start codons in transcripts derived from cryptic iTSSs but not in full-length mRNAs. We observed the start-codon signature in the *set2∆* strain but not in the wild type strain (Fig. 4C). This is particularly striking as even in the *set2∆* strain, the ribosome protection pattern is a composite of the translation signature of both canonical and iTSSs-derived transcripts. This result suggests that cryptic transcripts are not only associated to polyribosomes, but that ribosomes can identify new start codons as canonical ones. Our 5PSeq analysis of the RNA degradation sensitive transcripts (CUTs, up regulated in *rrp6∆*) revealed that those also could encode peptides (Fig. S8C-D). The first predicted ORFs downstream of the CUTs TSSs present a clear translation initiation signature and also a protection peak at 17 nt upstream of the stop codon (as expected from a terminating ribosome). This effect was especially clear in those CUTS not overlapping with canonical transcripts (*i.e.*, non iTSSs).

Finally, we analyzed whether our predicted truncated polypeptides matched acetylated N-termini of proteins using a recently published proteomics dataset as a reference (Varland et al. 2018). In the original study the authors identify 1056 canonical protein N-terminal sites in a wild-type strain using N-Terminal COFRADIC, which is a technique that maps modified N-termini of proteins on a global scale (2011). As chromatin-sensitive iTSS are expressed, even to a lower level, also in a wild type we were able to detect after proteomic reanalysis 7 iTSS derived polypeptides (Supplemental Table S3; see methods for details). Specifically we confirmed the expression of truncated proteins for SAS4, ORC1, SWC4, CNA1, NST1 and SMC5 (Fig 5 and S9) and the expression of an iTSS dependent peptide encoded in the 3’UTR of MON2 (Fig 5C). In addition, by comparing our 5’cap dataset with the one obtained by Doris *et al.* for *spt6-1004*, we can identify truncated transcripts previously shown by Western blot to produce also truncated proteins (Cheung et al. 2008).

**Figure 5.**
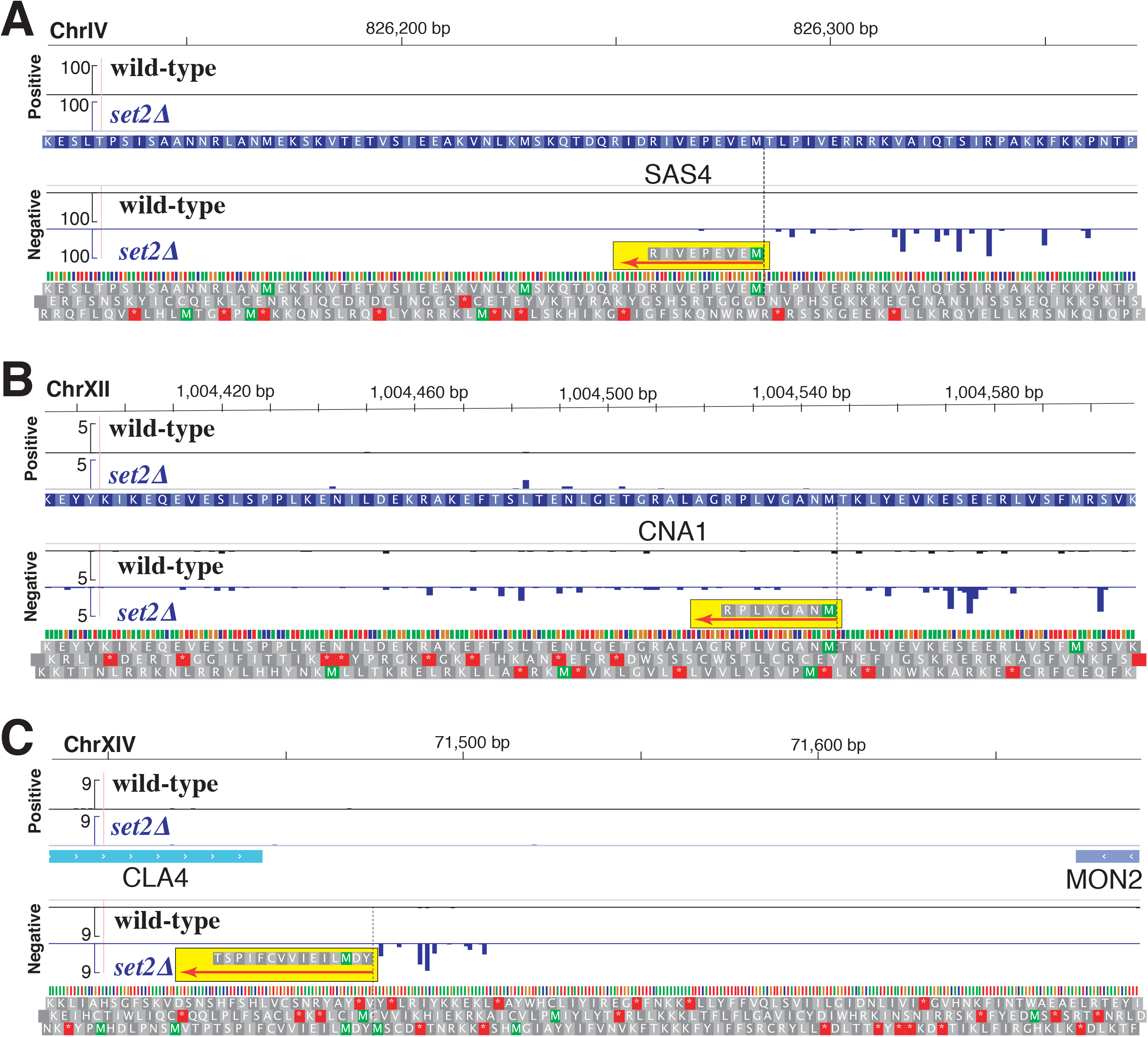
Chromatin-sensitive iTSSs encode peptides that can be detected by MS. Sequencing track display the 5’cap sequence Score (normalized counts) of collapsed replicates for wild-type (in black) and *∆set2* (in blue). Identified N-terminal peptides are highlighted in yellow and their orientations displayed using a red arrow. We display in grey the 3 potential translations of DNA in the same orientation of the detected peptide. A) Truncation of SAS4 (MEVEPEVIR). B) Truncation of CNA1 (MNAGVLPR). C) Chromatin-sensitive transcript encoding a peptide in the 3’UTR of MON2 (YDMLIEIVVCFIPST). N-terminal COFRADIC data from (Varland et al. 2018).

## Discussion

Here, we have shown that chromatin-sensitive cryptic promoters present multiple features similar to canonical gene-coding promoters. We focused on *set2∆-, rco1∆-* and *eaf3∆*-sensitive internal cryptic TSSs, and demonstrated that their DNA sequence, transcription directionality and chromatin organization are similar to those of canonical promoters (Fig. 1 and 2). This is in line with the characterization of cryptic promoters in the chaperone mutant *spt6-1004* that was published during the review of this manuscript (Doris et al. 2018). Our MNase footprint analysis showed that those promoters present a canonical nucleosome array organization and suggested that canonical TFs bind upstream of the iTSSs in the body of genes and are associated to the appearance of intergenic NFR (Fig. 2). Our observations are in agreement with recent reports that demonstrate how Gcn4 binds frequently in coding regions and can activate transcription from internal promoters (Rawal et al. 2018; Mittal et al. 2017). This suggests that a significant fraction of the cryptic promoters are in fact alternative promoters, whose expression under standard conditions is restricted by the chromatin organization or the absence of a particular transcription factor. Previous studies have shown that a significant number of Set2-repressed cryptic promoters can be regulated by carbon sources (Kim et al. 2016). Altogether, this suggests that our classification of cryptic and canonical promoters may be ingluenced by the environmental conditions under which cells are profiled.

To assess to what degree these cryptic iTSSs could represent *bona Bide* alternative transcript isoforms, we investigated their full boundaries (Fig 3). Using our previously developed TIF-seq approach, we identified that most of them employ the canonical polyadenylation sites used by full-length isoforms. Previous work from the Jacquier lab has shown that, by studying the double mutant *upf1∆ set2∆*, a proportion of internal cryptic transcripts are degraded by Non-sense Mediated Decay (NMD) (Malabat et al. 2015). Here we focused on the molecules that are present at a detectable level with active NMD pathway and thus more likely to have a post-transcriptional effect. We found that, even in a wild type strain, chromatin-sensitive iTSSs are typically associated to polyribosome fractions. To further dissect if these short isoforms are not only bound to polyribosomes, but actually translated, we applied an optimized version of our 5PSeq approach. We identified that the first methionine in the truncated transcripts presents a ribosome protection signature characteristic of translation start sites. In contrast, this signal is not detected in the wild type strain, in which truncated isoforms are expressed at low levels. Finally we reanalyzed a proteomics dataset of N-Terminally acetylated protein N-termini expressed in Wild type cells (Varland et al. 2018) and we found newly truncated protein isoforms based on our isoform predictions. Our observation extends on previous observations from our group and others showing that variations in the transcripts’ 5’ boundaries potentially leading to truncated proteins are common in yeast. Our results are in line with seminal work from the Winston group showing that the histone chaperone mutant *spt6-1004* can produce truncated proteins as analyzed by Western blot (Cheung et al. 2008). These variations may be environmentally regulated or occur simultaneously in a apparently homogenous population of cells (Pelechano et al. 2013; Fournier et al. 2012; Lycette et al. 2016; Carlson et al. 1983; Varland et al. 2018).

N-terminal proteomics approaches showed that downstream in-frame methionines often define alternative amino termini in the budding yeast proteome (Fournier et al. 2012; Lycette et al. 2016; Varland et al. 2018). These alternative proteoforms can be detected even in standard laboratory conditions suggesting that their expression coexist with the full-length proteoforms. However, most studies focused their analysis on the transcripts’ first 100 nucleotides, and thus did not investigate the downstream truncations that were commonly disregarded as cryptic transcripts. A similar phenomenon has been described in human cells, where alternative N-terminal proteoforms can lead to different protein stability (Gawron et al. 2016; Na et al. 2018). Regardless of their origin, it is clear that truncated proteins can have significant phenotypical impacts such as changes in protein localization (Carlson et al. 1983) or may even act as dominant-negative factors opposing the function of the full-length protein (Ungewitter and Scrable 2010). Our results also reveal that a fraction of CUTs have also the potential of encoding peptides. This is particularly intriguing, as CUTs are naturally unstable and thus the potential production of peptides would be also transient. In the future, further characterizing the abundance and functionality of alternative proteoforms derived from previously considered “cryptic” transcripts will be extremely valuable.

Although we focused our study on budding yeast, our conclusion that chromatin-sensitive cryptic iTSSs may act as alternative canonical TSSs have further implications. In mammals, alternative transcription start and termination sites, rather than alternative splicing, accounts for the majority of isoform differences across tissues (Reyes and Huber 2017). This highlights the importance of TSS selection in the definition of the transcriptome. It has been recently reported that the treatment of human cancer cell lines with DNA methyltransferase and histone deacetylase inhibitors (DNMTi and HDACi, respectively) results in the appearance of thousands of unannotated TSSs (TINATs) (Brocks et al. 2017). TINATs frequently splice into coding-protein exons and, in some cases, are associated with polyribosomes. Thus disruption of the epigenome by the DNMTi and HDACi treatments leads to the expression of cryptic TSSs similar to the chromatin-sensitive iTSSs defined here, both in terms of biogenesis and potential post-transcriptional consequences. This suggests that the expression of cryptic TSSs is likely to be evolutionary conserved and a source of alternative (functional or aberrant) proteoforms that should be further investigated. The study of chromatin-sensitive cryptic promoter regulation will help to better distinguish spurious transcripts from those functionally relevant although only expressed in a subpopulation of cells or under specific environmental conditions.

## Methods

### Cell growth

All *Saccharomyces cerevisiae* strains used in this study were derived from BY4741 (*MATa his3Δ1 leu2Δ0 met15Δ0 ura3Δ0*). BY4741, *rrp6∆* (*rrp6*::kanMX4)*, set2∆* (*set2*::kanMX4)*, rco1∆* (*rco1*::kanMX4) and *eaf3∆* (*eaf3*::kanMX4) were obtained from Euroscarf. *set1∆* (*set1*::kanMX4) was generated using standard yeast chemical transformation as previously described (Chabbert et al. 2015). Cells were grown in YPD (1% yeast extract, 2% peptone, 2% glucose and 40 mg/L adenine) and harvested at OD600 ~1. For 5PSeq start codon identification, cells were shifted for 5 minutes to YP media without glucose (1% yeast extract, 2% peptone) prior to harvesting. For 5PSeq in presence of cycloheximide, 0.1 mg/ml final cycloheximide was added for 10 minutes prior to harvesting. Total RNA was phenol extracted using standard methods and contaminant DNA was removed by DNase treatment (Turbo DNA-free kit, Ambion) (Pelechano et al. 2012).

### 5’cap library preparation

Identification of 5’capped mRNAs was performed as previously described (Pelechano et al. 2016). In brief, 10µg total RNA was treated with Calf intestinal alkaline phosphatase (NEB) to remove 5′P from fragmented and non-capped molecules. After purification, mRNA caps were removed using 3.75 units of Cap-Clip (Biozyme) exposing a 5’P in those molecules previously capped. Samples were ligated overnight at 16ºC with a DNA / RNA oligo (rP5 _ RND: TTTCCCTACACGACGCTCTTCCGATrCrUrNrNrNrNrNrNrNrN) using T4 RNA ligase 1 (New England Biolabs). RNA integrity after ligation was assayed by agarose gel electrophoresis and poly(A)RNA was purified using oligo dT magnetic beads. After this, ligated mRNA was fragmented at 80ºC for 5 min in the presence of RNA fragmentation buffer (40 mM Tris-acetate, pH 8.1, 100 mM KOAc, 30 mM MgOAc). Ligated RNA was subjected to reverse transcription using random hexamers with SuperScript II (Life Technologies) with the following program: 10 min at 25ºC, 50 min at 42ºC and heat inactivated for 15 min at 72ºC. Second strand cDNA synthesis was performed by a single PCR cycle (1 min at 98ºC; 2 min at 50ºC and 15 min at 72ºC) using Phusion High-Fidelity PCR Master Mix (New England Biolabs). A biotinylated oligo (Bio NotI-P 5 -P ET: [Btn]TATAGCGGCCGCAATGATACGGCGACCACCGAG ATCTACACTCTTTCCCTA CACGACGCTCTTCCGATCT) was added during the generation of the second cDNA strand. Double stranded cDNA was purified using Ampure XP (Beckman Coulter) or HighPrep (Magbio) beads. After the samples were bound to streptavidin coated magnetic beads (M-280 Dynabeads, Life Technologies) and subjected to standard Illumina end-repair, dA addition and adapter ligation was performed as previously described (Pelechano et al. 2016). Libraries were enriched by PCR and sequenced in an Illumina HiSeq 2000 instrument.

### TIF-seq sequencing

TIF-seq libraries were performed as described in (Pelechano et al. 2013) using 60 µg of DNA-free total RNA as input. In brief, 5’ non-capped molecules were dephosphorylated using 6 units of Shrimp Alkaline Phosphatase (Fermentas). RNA was phenol purified, and the 5’P of capped molecules was exposed by treatment with 5 units of Tobacco Acid Pyrophosphatase (Epicentre). RNA samples were ligated with the TIF-seq DNA/ RNA 5oligo cap using T4 RNA ligase 1 (NEB). Full-length cDNA (FlcDNA) was generated with SuperScript III reverse transcriptase and amplified by PCR with HF Phusion MasterMix (Finnzymes).

FlcDNA was digested with NotI (NEB) to generate cohesive ends. Samples were subjected to intramolecular ligation using T4DNA ligase. TIF-seq chimeras were controlled mixing 2 aliquots of differentially barcoded FlcDNA during the ligation, as described in the original TIF-seq manuscript. Non-circularized molecules were degraded using Exonuclease III and Exonuclease I (NEB). Circularized FlcDNAs was fragmented by sonication using a Covaris S220 (4 min, 20% Duty Cycle, Intensity 5, 200 cycles/burst). Fragmented DNA was purified, and biotin-containing fragments were captured with Streptavidin-conjugated Dynabeads M-280 (Invitrogen). Forked barcoded adapters were added using the standard Illumina DNA-seq library generation protocols. Libraries were enriched by 20 cycles of PCR Phusion polymerase (Finnzymes). 300 bp libraries were isolated using e-Gel 2% SizeSelect (Invitrogen) and sequenced in an Illumina HiSeq 2000 instrument (105 paired-end sequencing).

### Polyribosome fractionation

100 mL of *S. cerevisiae* cells at OD_600_ ~1 were treated with cycloheximide for 5 minutes (100 µg/mL, final concentration), harvested by centrifugation and transferred to ice. Pellets were washed with ice-cold lysis buffer and resuspended in 700µL lysis buffer. Lysis buffer contains 20 mM Tris-HCl, pH 8, 140 mM KCl, 5 mM MgCl_2_, 0.5 mM DTT, 1% Triton X-100, 100 µg/ mL cycloheximide, 500 µg/mL heparin and complete EDTA-free protease inhibitor (1 tablet per 10 mL, Sigma Aldrich). For cell lysis, samples were transferred to pre-cooled 1.5mL screw-tubes with 300 µL glass beads and supplemented with 100 units of RNAse inhibitor (RNAsin plus, Promega). Cells were lysed using a FastPrep-24 shaker (6.0m/s for 15 seconds, MP biomedicals). Supernatant was recovered after 5 minutes centrifugation at 2300g, and cleared with an additional centrifugation at 5900g. Extracts were supplemented with glycerol (5% final v/v) and stored at −70C. 10-50% sucrose gradients were prepared with a Gradient Master BIOCOMP (Nycomed Pharma). Sucrose solution contains 20 mM Tris-HCl, pH 8, 140 mM KCl, 5 mM MgCl_2_, 0.5 mM DTT, 100 µg/mL cycloheximide and sucrose (from 10 to 50 %). Cleared cell extracts were ultracentifuged at 34400 rpm for 2 hours 40 minutes at 4C using a C-1000 XP centrifuge with SW40 rotor (Beckman Coulter). Gradient UV absorption at 254 nm was measured and selected fractions were selected for 5’cap library preparation (5µg purified RNA per sample). Polyribosome fraction (*i.e.*, 2n+) was compared with the total extract prior to fractionation).

### 5PSeq

5PSeq libraries were prepared as previously described (Pelechano et al. 2015; 2016). 5PSeq protocol is the same as the one described for 5’cap sequencing (see above) with variations only for the RNA ligation and rRNA depletion. Specifically, 6 µg of total RNA were directly ligated with a DNA/RNA oligo (rP5_RND). In that way only molecules with a 5’P in the original sample are ligated. Ribosomal RNAs were depleted using Ribo-Zero Magnetic Gold Kit (Illumina). Samples were sequenced in an Illumina NextSeq 500 instrument.

### Bioinformatic analysis

For 5’ cap sequencing reads, random barcodes were first extracted and added to the reads name. The reads were aligned to yeast genome (S. cerevisiae genome (SGD R64-1-1; sacCer3) with Novoalign (http://www.novocraft.com) using default setting. A customized script adapted from UMI-tools (Smith et al. 2017) was used for removing PCR duplicates (Supplemental Code S1). Specifically we allowed 1 bp shifting at the beginning of 5’ ends. CAGEr was employed for clustering the 5’ cap TSSs of BY4741 wild-type strain and the mutants (Haberle et al. 2015). TSS counts in different samples were normalized to match a common reference power-law distribution. Low-fidelity tags supported by less than 2 normalized counts in all samples were filtered out before clustering. In each sample, neighboring tags within 20 bp were spatially clustered into larger tag clusters. If the tag clusters were within 10 bp apart, they were aggregated together into non-overlapping consensus clusters across all samples. The raw expression counts of the consensus clusters were further exported to the DESeq2 (Love et al. 2014) for differential expression analysis, comparing between mutants and wild-type strain. Polyribosome derived 5’ cap sequencing reads were assigned to the consensus clusters by featureCounts (Liao et al. 2014), with read counting based on the 5’ most base. Differential expression analysis of polyribosome fractionation against total extract was performed using DESeq2.

Bar-ChIP sequencing data were processed as described previously (Chabbert et al. 2015).

TIF-seq sequencing data were processed as described previously (Pelechano et al. 2013). In general, all reads were first de-multiplexed and random barcodes were extracted. Pairs of transcript 5’ and 3’ end reads were mapped to yeast genome (S. cerevisiae genome (SGD R64-1-1; sacCer3) with Novoalign (http://www.novocraft.com) using default setting separately. Only transcripts with both ends mapped in same chromosome a length ranging from 40 to 5,000bp were used for further analysis.

5PSeq reads were mapped to the *S. cerevisiae* (genome R64-1-1) using STAR 2.5.3a (Dobin et al. 2013) with default parameters except AlignIntronMax (2500). PCR duplicates were removed as described for 5’ cap sequencing. Reads were aligned to either the start codon, or the first in-frame methionine downstream of *set2∆-*specific iTSS.

We analyzed the MS raw data from Varland et. al. 2018 (PRIDE: PXD004326) including our additional predictions. MS/MS peak lists were searched essentially as described in Varland et. al. 2018 using the Sequest database (Thermo Scientific). Spectral searches were performed using the UniProtKB Saccharomyces cerevisiae database (version 2018_08) supplemented with the putative truncated proteins encoded by in-frame methionines downstream of iTSS. To maximize our ability to detect iTSS derived N-terminal peptides expressed also in the wild-type strain, we relaxed the stringency of the iTSS selection to p-adjusted <0.05. 13C2D3-acetylation of lysine side-chains, carbamidomethylation of cysteine and methionine oxidation to methionine-sulfoxide were set as fixed modifications. 13C2D3-acetylation, acetylation of protein N termini and pyroglutamate formation of N-terminal glutamine were set as a variable modification. Mass tolerances on precursor ions were set to 10 ppm and on fragment ions to 0.5 Da. The estimated false discovery rate by searching decoy databases were below 1%. Similar results were obtained using the Mascot search database (Version 2.5, Matrix Science).

### Data access

All raw and processed sequencing data generated in this study have been submitted to the NCBI Gene Expression Omnibus (GEO; http://www.ncbi.nlm.nih.gov/geo/) under accession numbers: GSE119114, GSE119160, GSE118758, GSE119134 and GSE128599.

## Acknowledgments

We want to thank all members of Steinmetz, Pelechano and Wei laboratories for discussion. We thank Petra Jakob, Manu Tekkedil and Sandra Clauder-Münster for technical assistance. We thank Bruno Galy for his help with polyribosome fractionation. This study was technically supported by the European Molecular Biology Laboratory (EMBL) Genomics Core Facility, the Science for Life Laboratory (Sweden) and computational resources provided by SNIC through Uppsala Multidisciplinary Center for Advanced Computational Science (UPPMAX). This study was financially supported by the National Key R&D Program of China (2017YFC0908405) and National Natural Science Foundation of China (Grant No: 81870187) to W.W.; the US National Institutes of Health (NIH grant P01 HG000205), Deutsche Forschungsgemeinschaft (1422/4-1) and a European Research Council Advanced Investigator Grant to L.M.S.; and by the Swedish Research Council (VR 2016-01842), a Wallenberg Academy Fellowship (KAW 2016.0123), the Swedish Foundations’ Starting Grant (Ragnar Söderberg Foundation) and Karolinska Institutet (SciLifeLab Fellowship, SFO and KI funds) to V.P.; V.P and W.W. acknowledge the support from a Joint China-Sweden mobility grant from STINT (CH2018-7750) and the National Natural Science Foundation of China (NSFC) respectively; B.P.H. and S.H.A were supported by a fellowship from the EMBL Interdisciplinary Postdoctoral (EIPOD) programme under Marie Sklodowska-Curie Actions COFUND (grant number 291772). Y.Z. is funded by a fellowship from the China Scholarship Council. C.D.C was supported by a PhD fellowship from the Boehringer Ingelheim Fonds.

## Additional information

### Author contribution

Conceptualization: VP, WW & LSM; Acquisition of data and interpretation: VP, BPH, YZ, YPS, CDC & SHA; Computational analysis and interpretation: WW, VP, JW, IP & CDC; Writing – original draft preparation: VP & WW; Writing – review & editing: VP, WW, BPH, JW, YZ, YPS, CDC, SHA, IP & LSM; Supervision: VP & LSM; Funding acquisition: LSM, VP & WW.

