## Supplementary text and figures for "Chromatin-sensitive cryptic promoters encode alternative protein isoforms in yeast"

### Supplemental information

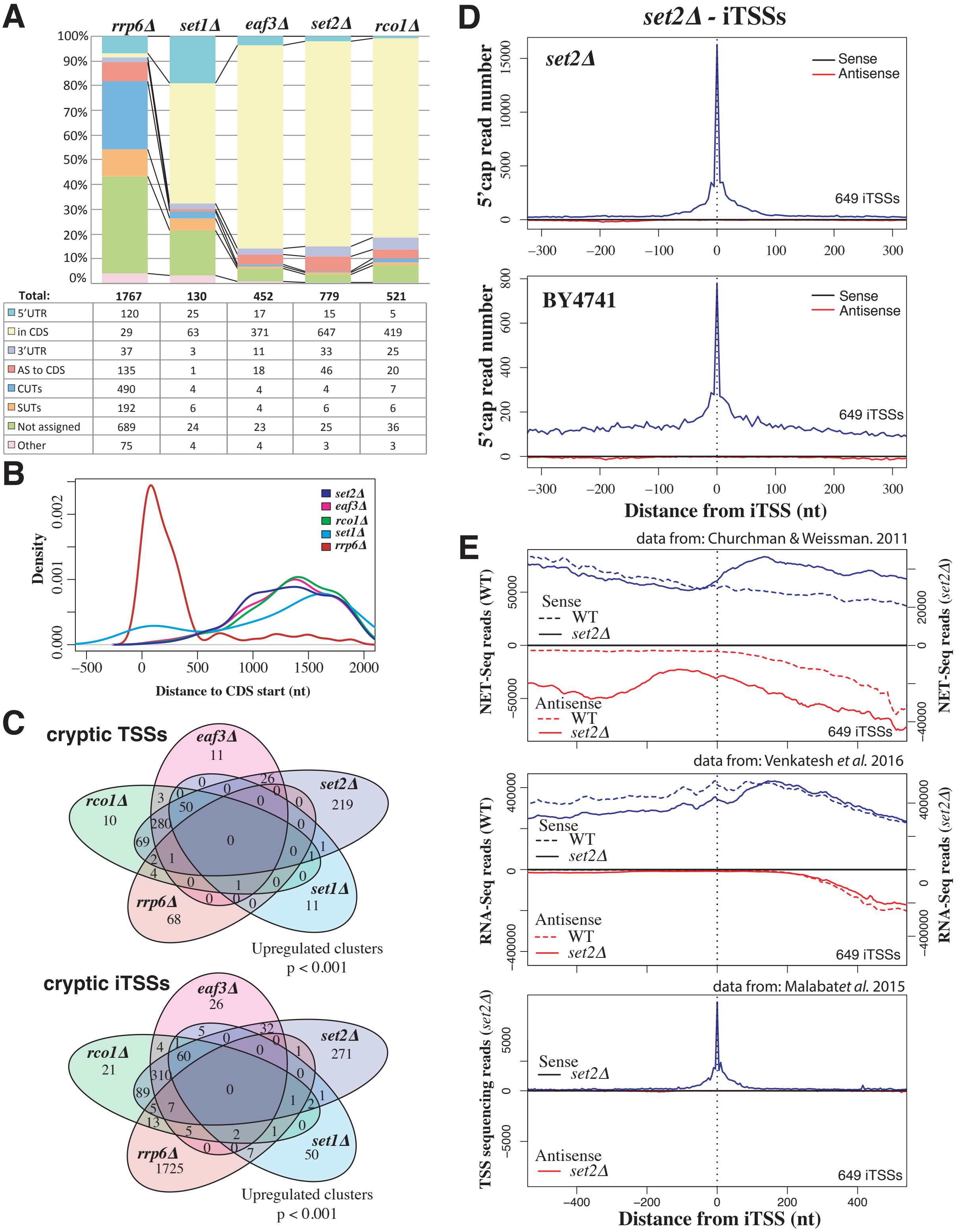


| Total: | 1767 | 130 | 452 | 779 | 521 |
|---|---|---|---|---|---|
| 5'UTR | 120 | 25 | 17 | 15 | 5 |
| in CDS | 29 | 63 | 371 | 647 | 419 |
| 3'UTR | 37 | 3 | 11 | 33 | 25 |
| AS to CDS | 135 | 1 | 18 | 46 | 20 |
| CUTs | 490 | 4 | 4 | 4 | 7 |
| SUTs | 192 | 6 | 4 | 6 | 6 |
| Not assigned | 689 | 24 | 23 | 25 | 36 |
| Other | 75 | 4 | 4 | 3 | 3 |

**Supplemental Figure S1.** Relationship between TSSs across studied strains. (A) Classification of differentially expressed TSSs in respect to annotated features. Annotation of SUTs (Stable Unannotated Transcripts), CUTS and UTR lengths are from (Xu et al. 2009). (B) Distribution of differentially expressed TSSs with respect to annotated start codons of ORFs. (C) Relationship between TSSs identified across strains. Analysis is shown for all TSSs (upper part) or those occurring inside coding regions in the same orientation (iTSSs, bottom part). (D) 5'cap sequencing reads with respect to the identified *set2Δ*-sensitive cryptic iTSSs. Reads in sense orientation (respect to the overlapping ORF) are depicted in blue, and antisense 5'cap reads in red. *set2Δ*-sensitive transcript are very apparent in *set2Δ*, but can be also observed at lower expression in the wild-type strain (BY4741). Please note that in the BY4741 the height of the iTSS peak in respect to the background sense transcription is much lower than in the *set2Δ*. (E) Comparison of *set2Δ*-sensitive cryptic iTSSs in alternative datasets. NET-seq (Churchman and Weissman 2012) and RNA-seq data (Venkatesh et al. 2016) depict the coverage for the same regions in a wild-type strain (dashed lines) and for *set2Δ* (solid lines). Comparison with an independent TSS sequencing dataset (Malabat et al. 2015) is shown as in C. It can be observed that nascent transcription arises bidirectionally from the cryptic promoters (specially in *set2Δ* NET-seq data). But that the stable detected cryptic transcripts (5'cap, RNA-seq or TSS sequencing) are directional.

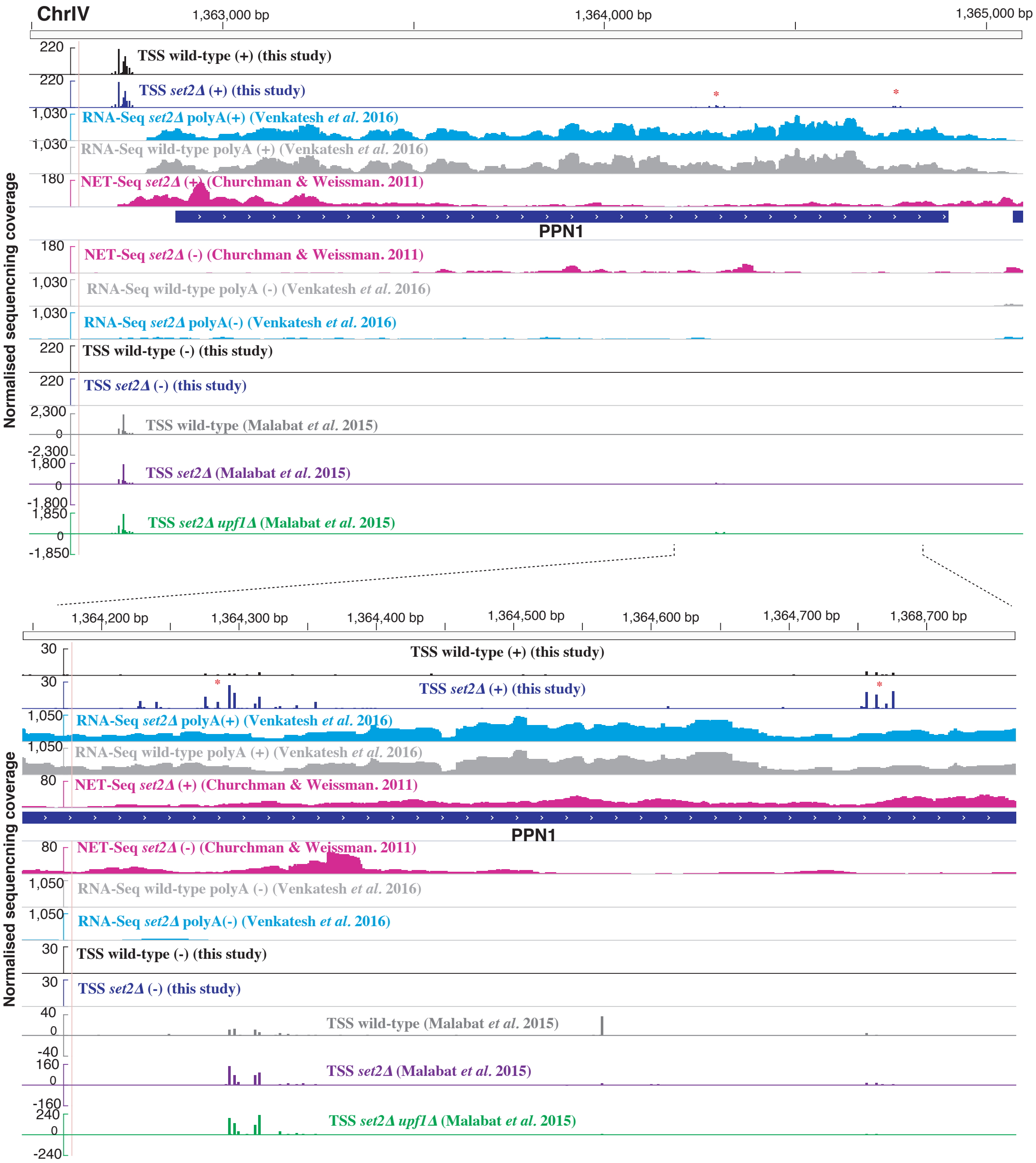


**Supplemental Figure S2.** Representative sequencing track comparing the dataset obtained in this study (as in Fig 1A) with alternative datasets from NET-seq (Churchman and Weissman 2012), RNA-seq (Venkatesh et al. 2016) and TSS sequencing (Malabat et al. 2015).

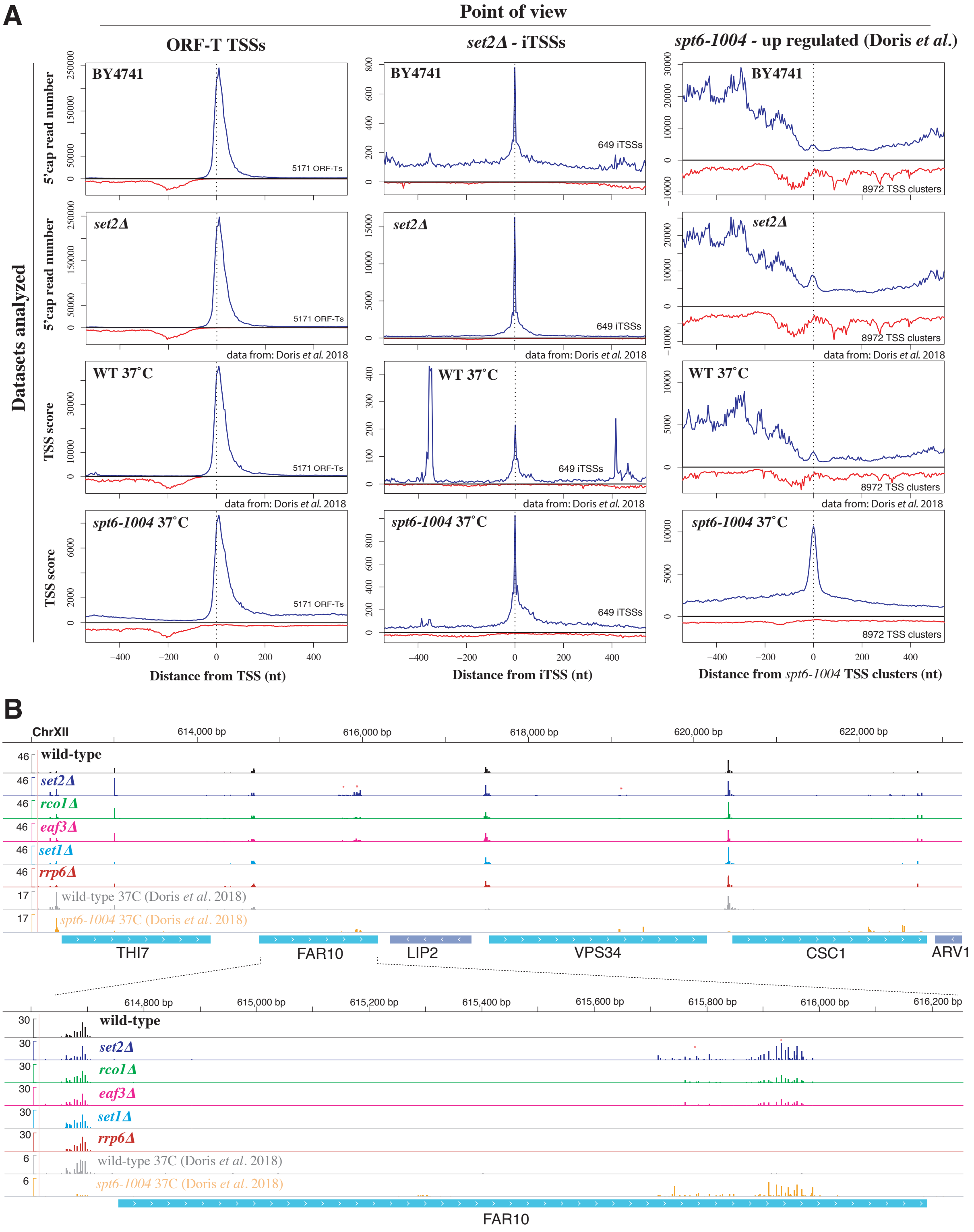


**Supplemental Figure S3.** Comparison between *spt6-1004* and chromatin-sensitive iTSSs. (A) Comparison of 5´cap coverage in respect to canonical ORF-T promoters, *Δset2* iTSSs and *spt6-1004* dependent promoters (Log$_2$ fold-change >0 and p-adjusted <0.1, as defined in Supplemental table S1 from (Doris et al. 2018)). Reads in sense orientation (respect to the overlapping ORF) are depicted in blue, and antisense 5'cap reads in red. (B) Representative sequencing track comparing the dataset obtained in this study with *spt6-1004* transcription start sequencing datasets (Doris et al. 2018).

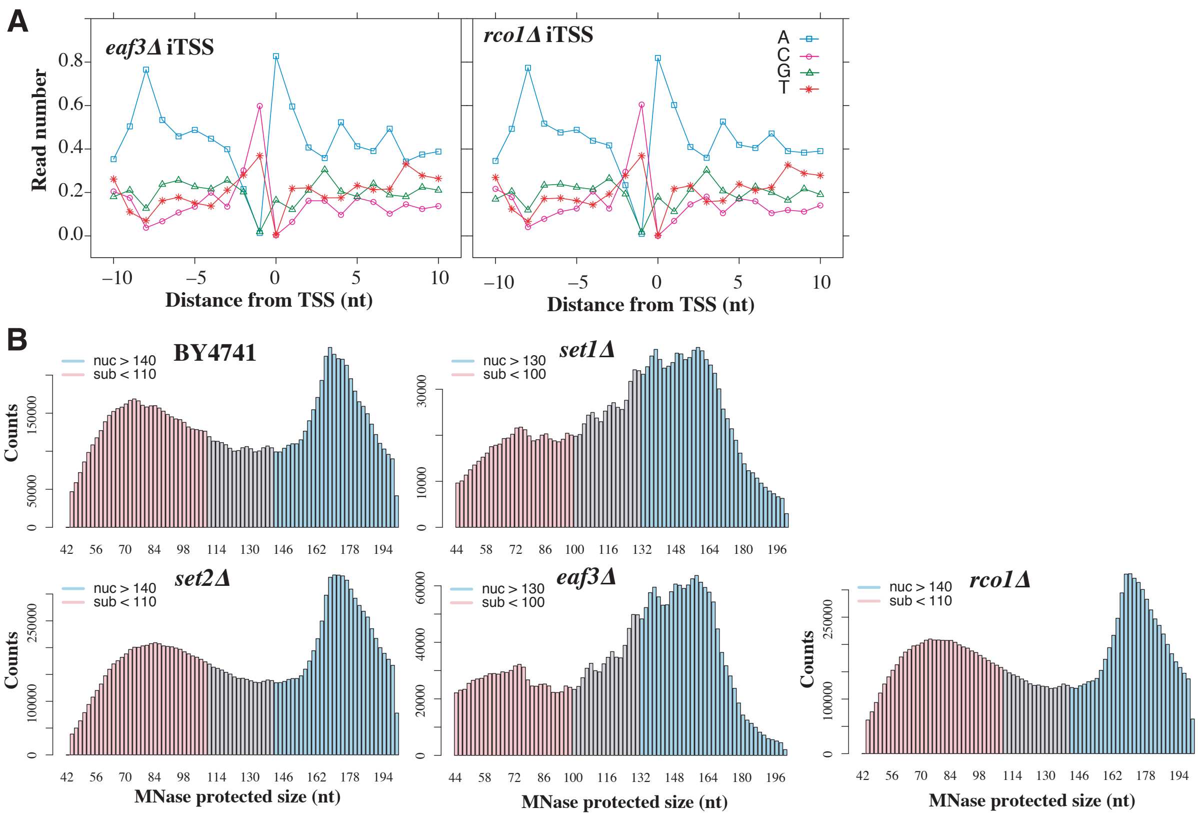


**Supplemental Figure S4.** Sequence preference of iTSSs and size of MNase fragments. (A) Sequence preference of *eaf3Δ* and *rco1Δ* iTSSs. (C) Size distribution of analyzed MNase protected fragments. Selected nucleosome protection fragments (nuc) and sub-nucleosomal fragments (sub) are depicted in blue and pink respectively. Chromatin data is reanalysed from (Chabbert et al. 2015).

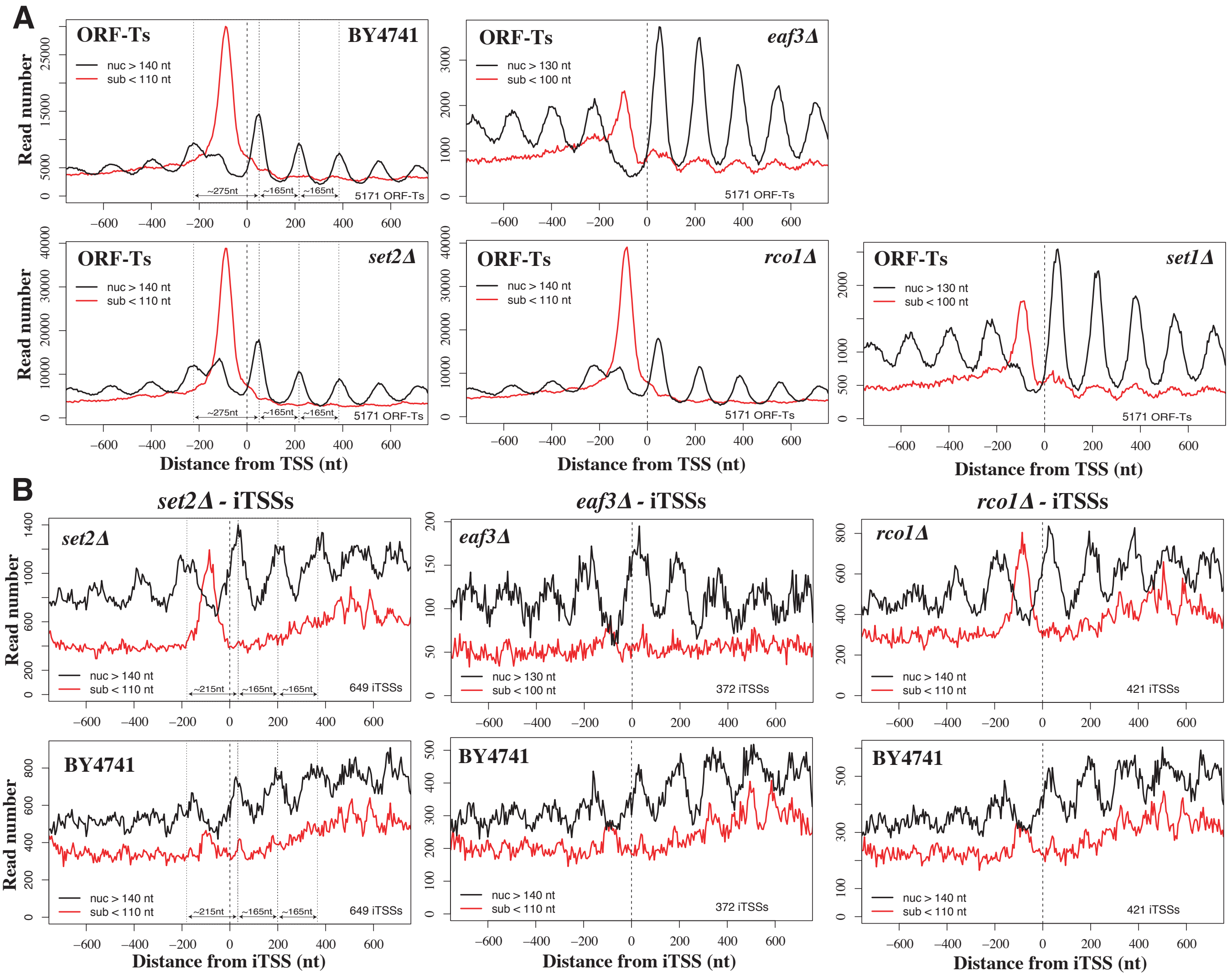


**Supplemental Figure S5.** MNase protection pattern of iTSSs. (A) MNase protection pattern for canonical ORF-T TSSs in different strains. MNase fragments are distributed in nucleosome protection fragments (nuc) and sub-nucleosomal ones (sub) according to their length. (B) MNase protection pattern for cryptic up-regulated iTSSs for *set2Δ*, *rco1Δ* and *eaf3Δ*. Chromatin data is reanalysed from (Chabbert et al. 2015).

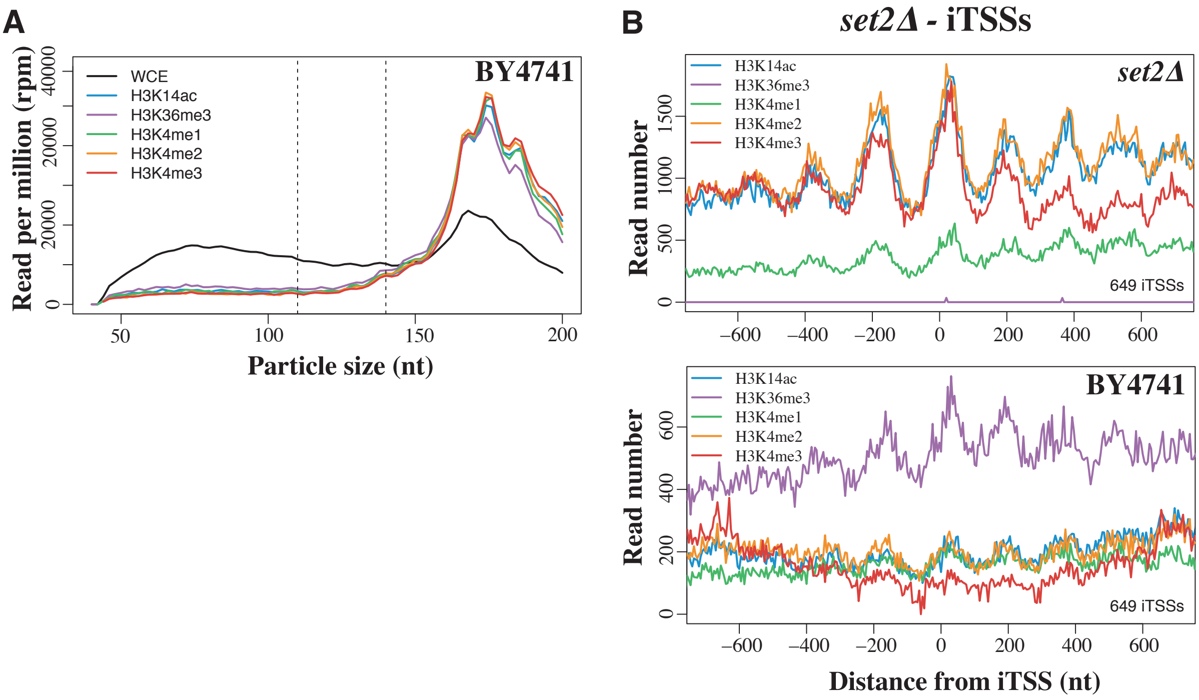


**Supplemental Figure S6.** MNase protected fragments versus histone association (A) Length distribution of MNase protected fragments prior (in black) or after histone immunoprecipitation. Subnucleosomal fragments are depleted when histones are immunoprecipitated. (B) Histone marks distribution profiles around *set2Δ* iTSSs. H3K4me3 (in red) is high in the "cryptic +1 nucleosome" and decreases towards the body of the cryptic transcript. Chromatin data is reanalysed from (Chabbert et al. 2015).

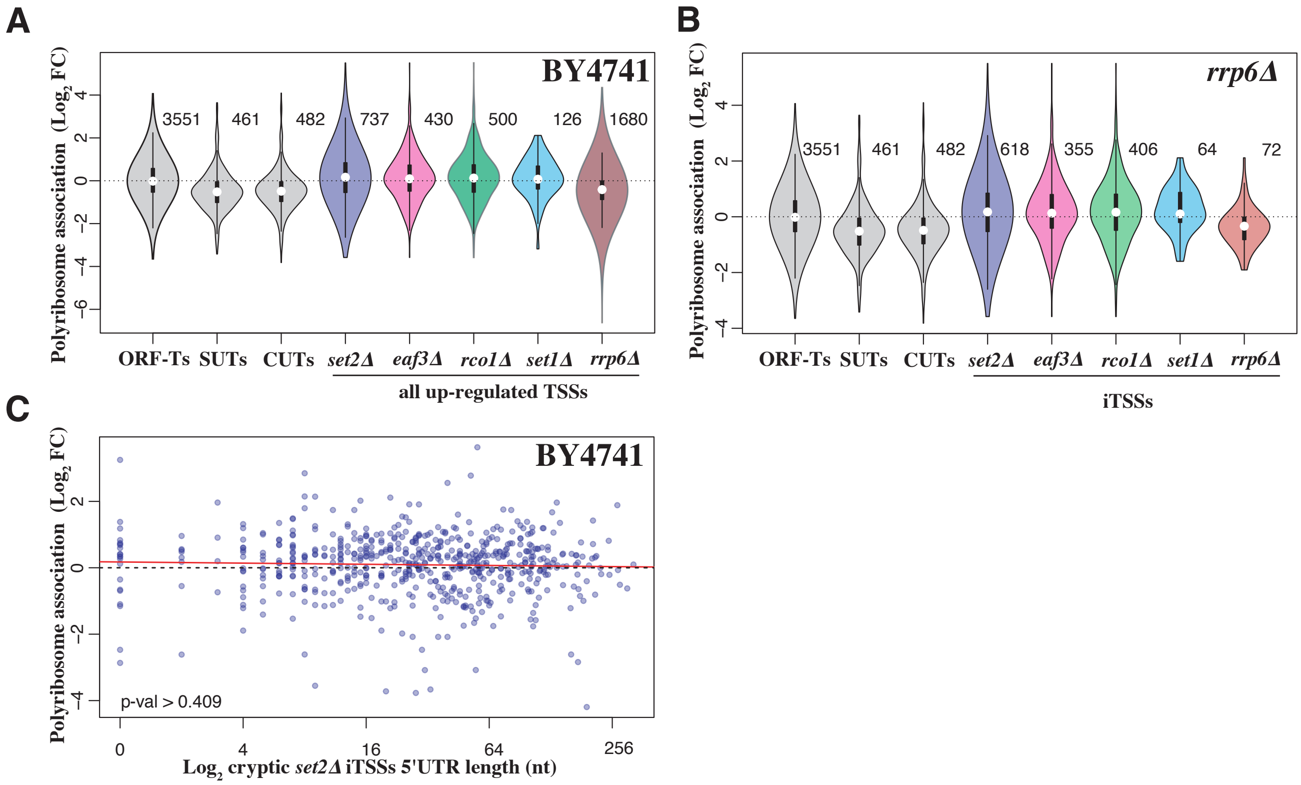


**Supplemental Figure S7.** Polyribosome association of cryptic transcript. (A) Relative association with polyribosome fraction after sucrose fractionation versus total extract in a wild type strain (as in Fig. 4A). All cryptic up-regulated TSSs (independent of their location) are shown. Analyzed events (present at a sufficient level in the wild-type strain) are indicated to the right of each plot. (B) Relative association with polyribosome fraction after sucrose fractionation versus total extract in a *rrp6Δ* strain. Analyzed events (present at a sufficient level in the wild-type strain) are indicated to the right of each plot. (C) Distribution of the length between the *set2Δ*-sensitive cryptic iTSSs and the first in frame ATG (cryptic 5'UTR, in x-axes) versus the relative polyribosome association (y-axis, as in Fig 4A). No significant association was detected.

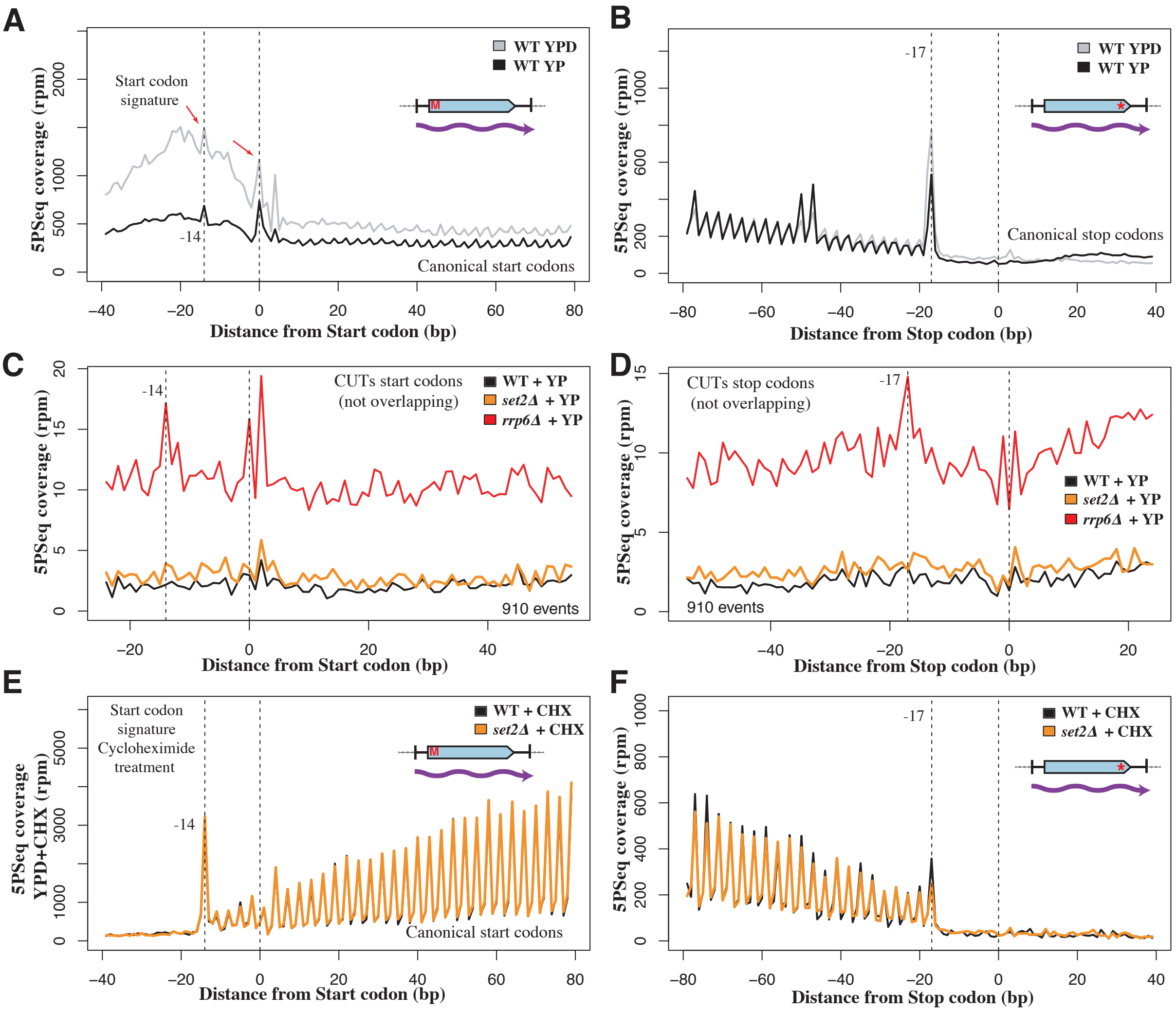


**Supplemental Figure S8.** 5PSeq protection pattern of canonical mRNAs and CUTs. (A-B) Comparison of 5PSeq start-codon associated signature before (grey) and after glucose depletion (black) for coding genes. Glucose depletion leads to a decrease in the coverage around the 5' region of the genes (left side), and only to a subtle increase in pause at the stop codon (right side). To decrease the effect of potential outliers, we assigned a value corresponding to the 95th percentile to values that were over this threshold at each distance from the start codon. (C-D) 5PSeq start-codon associated signature after glucose depletion for different mutants. The first putatively encoded ORF downstream of each RNA degradation sensitive TSSs (CUTs) was used to perform a metagene analysis. Only those predicted ORFs not overlapping with ORF-Ts and other genomic features (snRNAs, tRNAS...) were considered. A clear protection pattern can be observed for the start and the stop codon of the putative ORFs present in the CUTs (in red). To decrease the effect of outliers, we assigned a value corresponding to the 80th percentile to values that were over this threshold at each distance from the start codon. (E-F) Effect of cycloheximide treatment in 5PSeq profiles. It leads to an increase at the position -14 nt from the start site and also to an increase ribosome stalling thought the gene body. As in A.

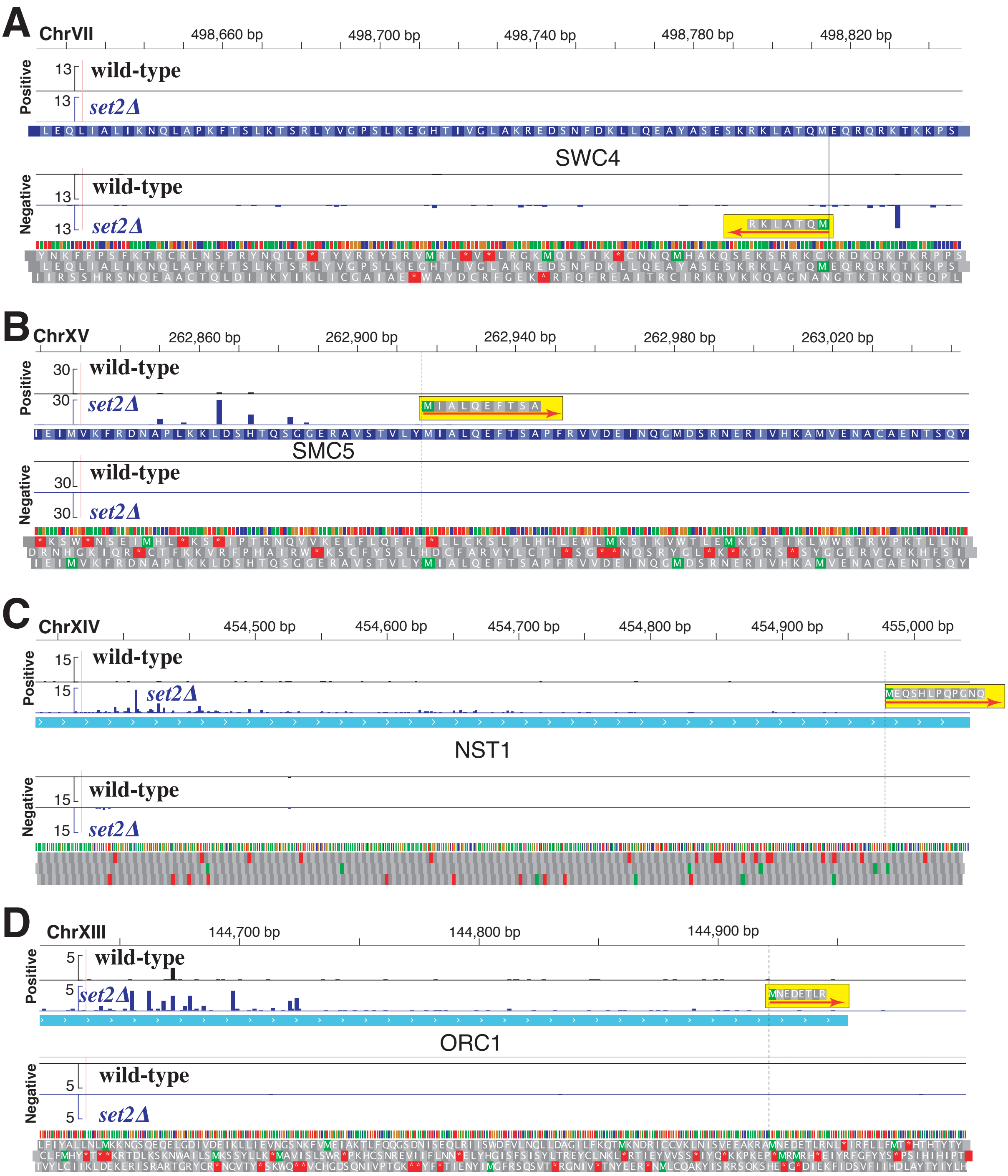


**Supplemental Figure S9.** Additional chromatin-sensitive iTSSs encoded peptides identified by MS. (A) Truncation of SWC4 (MQTALKR). (B) Truncation of SMC5 (MIALQEFTSA). (C) Truncation of NST1(MEQSHLPQPGNQ) (D) Truncation of ORC1 (MNEDETLR). Please note that in the case of NST1 and ORC1 the peptide is encoded downstream of the first Methionine. N-terminal COFRADIC data from (Varland *et al.* 2018)

**Supplemental Table S1.** Text file with all identified TSSs and their fold change expression versus wild-type as measured by 5'cap sequencing.

**Supplemental Table S2.** Text file with differential expression for the presence of 5' cap molecules in polyribosome fractions with respect the total extract. Experiment were performed in the wild-type strain (BY4741).

**Supplemental Table S3.** MS evidence for reanalyzed N-terminal proteomics dataset.

**Supplemental Code S1.** Modified UMI-tools python script.

**Used datasets:**

| Data | Accession | Origin |
|---|---|---|
| 5'cap and polyribosomes | GEO: GSE119114 | This study |
| TIF-seq set2Δ | GEO: GSE119160 | This study |
| 5PSeq: wild-type and set2Δ | GEO: GSE118758 | This study |
| 5PSeq: YPD and Cyclohemide | GEO: GSE128599 | This study |
| 5PSeq: rrp6Δ and polyribosomes | GEO: GSE119134 | This study |
| Bar-ChIP | ENA: ERP007035 | (Chabbert et al. 2015) |
| RNA-seq: wild-type and set2Δ | SRA: SRP089706 | (Venkatesh et al. 2016) |
| NET-seq: wild-type and set2Δ | GEO: GSE25107 | (Churchman and Weissman 2012) |
| TSS sequencing | GEO: GSE64139 | (Malabat et al. 2015) |
| TSS sequencing ***spt6-1004*** | GEO: GSE115775 | (Doris et al. 2018) |
| N-terminal COFRADIC | PRIDE: PXD004326 | (Malabat et al. 2015) |
