## Supplemental code for "Chromatin-sensitive cryptic promoters encode alternative protein isoforms in yeast": dedup.py

```
'''
Modified based on UMI-tools source codes
Compatible in python 2.7
'''
import os
import sys
import pysam
import collections
import itertools
import random
import logging
import re
from functools import partial
import numpy as np
import optparse
import textwrap
import copy
import time
import gzip
import inspect
import uuid
import tempfile
#import Utilities as U

global_id = uuid.uuid4()
global_benchmark = collections.defaultdict(int)

def hamming_distance(s1, s2):
    assert len(s1) == len(s2)
    return sum(ch1 != ch2 for ch1, ch2 in zip(s1, s2))

def breadth_first_search(node, adj_list):
    searched = set()
    queue = set()
    queue.update((node,))
    searched.update((node,))

    while len(queue) > 0:
        node = queue.pop()
        for next_node in adj_list[node]:
            if next_node not in searched:
                queue.update((next_node,))
                searched.update((next_node,))

    return searched

def get_substr_slices(umi_length, idx_size):
    '''
    Create slices to split a UMI into approximately equal size substrings
    Returns a list of tuples that can be passed to slice function
    '''
    cs, r = divmod(umi_length, idx_size)
    sub_sizes = [cs + 1] * r + [cs] * (idx_size - r)
    offset = 0
    slices = []
    for s in sub_sizes:
        slices.append((offset, offset + s))
        offset += s
    return slices

def build_substr_idx(umis, umi_length, min_edit):
    '''
    Build a dictionary of nearest neighbours using substrings, can be used
    to reduce the number of pairwise comparisons.
    '''
    substr_idx = collections.defaultdict(
        lambda: collections.defaultdict(set))
    slices = get_substr_slices(umi_length, min_edit + 1)
    for idx in slices:
        for u in umis:
            u_sub = u[slice(*idx)]
            substr_idx[idx][u_sub].add(u)
    return substr_idx

def iter_nearest_neighbours(umis, substr_idx):
    '''
    Added by Matt 06/05/17
    use substring dict to get (approximately) all the nearest neighbours to
    each in a set of umis.
    '''
    for u in umis:
        neighbours = set()
        for idx, substr_map in substr_idx.items():
            u_sub = u[slice(*idx)]
            neighbours = neighbours.union(substr_map[u_sub])
        neighbours.remove(u)
        for nbr in neighbours:
            yield u, nbr

def find_splice(cigar):
    '''Takes a cigar string and finds the first splice position as
    an offset from the start. To find the 5' end (read coords) of
    the junction for a reverse read, pass in the reversed cigar tuple'''

    offset = 0
    # a soft clip at the end of the read is taken as splicing
    # where as a soft clip at the start is not.
    if cigar[0][0] == 4:
        offset = cigar[0][1]
        cigar = cigar[1:]

    for op, bases in cigar:
        if op in (3, 4):
            # N or S: found the splice
            return offset
        elif op in (0, 2, 7, 8):
            # M, D, = or X: reference consuming
            offset += bases
        elif op in (1, 5, 6):
            # I, H, P: non-reference consuming
            continue
        else:
            raise ValueError("Bad Cigar operation: %i" % op)

    return False

def get_read_position(read, soft_clip_threshold):
    ''' get the read position (taking account of clipping) '''
    is_spliced = False

    if read.is_reverse:
        pos = read.aend
        if read.cigar[-1][0] == 4:
            pos = pos + read.cigar[-1][1]
        start = read.pos

        if ('N' in read.cigarstring or
            (read.cigar[0][0] == 4 and
             read.cigar[0][1] > soft_clip_threshold)):

            cigar = read.cigar[::-1]
            is_spliced = find_splice(cigar)
    else:
        pos = read.pos
        if read.cigar[0][0] == 4:
            pos = pos - read.cigar[0][1]
        start = pos

        if ('N' in read.cigarstring or
            (read.cigar[-1][0] == 4 and
             read.cigar[-1][1] > soft_clip_threshold)):
            is_spliced = find_splice(read.cigar)

    return start, pos, is_spliced

def get_barcode_read_id(read, cell_barcode=False, sep="_"):
    ''' extract the umi +/- cell barcode from the read id using the
    specified separator '''

    try:
        if cell_barcode:
            umi = read.qname.split(sep)[-1].encode('utf-8')
            cell = read.qname.split(sep)[-2].encode('utf-8')
        else:
            umi = read.qname.split(sep)[-1].encode('utf-8')
            cell = None

        return umi, cell

    except:
        raise ValueError(
            "Could not extract UMI +/- cell barcode from the read"
            "ID, please check UMI is encoded in the read name")

def remove_umis(adj_list, cluster, nodes):
    '''removes the specified nodes from the cluster and returns
    the remaining nodes '''

    # list incomprehension: for x in nodes: for node in adj_list[x]: yield node
    nodes_to_remove = set([node
                           for x in nodes
                           for node in adj_list[x]] + nodes)

    return cluster - nodes_to_remove

def get_average_umi_distance(umis):

    if len(umis) == 1:
        return -1

    dists = [hamming_distance(x, y) for
             x, y in itertools.combinations(umis, 2)]
    return float(sum(dists))/(len(dists))

def detect_bam_features(bamfile, n_entries=1000):
    ''' read the first n entries in the bam file and identify the tags
    available detecting multimapping '''

    inbam = pysam.Samfile(bamfile)
    inbam = inbam.fetch(until_eof=True)

    tags = ["NH", "X0", "XT"]
    available_tags = {x: 1 for x in tags}

    for n, read in enumerate(inbam):
        if n > n_entries:
            break

        if read.is_unmapped:
            continue

        else:
            for tag in tags:
                if not read.has_tag(tag):
                    available_tags[tag] = 0

    return available_tags

class UMIClusterer:
    '''A functor that clusters a dictionary of UMIs and their counts.
    The primary return value is either a list of representative UMIs
    or a list of lists where each inner list represents the contents of
    one cluster.

    Optionally:

      - identify the parent UMIs and return:
         - selected reads
         - umis
         - counts

    The initiation of the functor defines the methods:

      ** get_adj_list ** - returns the edges connecting the UMIs

      ** get_connected_components ** - returns clusters of connected components
                                       using the edges in the adjacency list

      ** get_groups ** - returns the groups of umis,
                         with the parent umi at position 0

    Note: The get_adj_list and connected_components methods are not required by
    all custering methods. Where there are not required, the methods return
    None or the input parameters.

    '''

    # "get_best" methods #

    def _get_best_min_account(self, cluster, adj_list, counts):
        ''' return the min UMI(s) need to account for cluster'''
        if len(cluster) == 1:
            return list(cluster)

        sorted_nodes = sorted(cluster, key=lambda x: counts[x],
                              reverse=True)

        for i in range(len(sorted_nodes) - 1):
            if len(remove_umis(adj_list, cluster, sorted_nodes[:i+1])) == 0:
                return sorted_nodes[:i+1]

    def _get_best_percentile(self, cluster, counts):
        ''' return all UMIs with counts >1% of the
        median counts in the cluster '''

        if len(cluster) == 1:
            return list(cluster)
        else:
            threshold = np.median(list(counts.values()))/100
            return [read for read in cluster if counts[read] > threshold]

    # "get_adj_list" methods #

    def _get_adj_list_adjacency(self, umis, counts, threshold):
        ''' identify all umis within hamming distance threshold'''

        adj_list = {umi: [] for umi in umis}
        if len(umis) > 25:
            umi_length = len(umis[0])
            substr_idx = build_substr_idx(umis, umi_length, threshold)
            iter_umi_pairs = iter_nearest_neighbours(umis, substr_idx)
        else:
            iter_umi_pairs = itertools.combinations(umis, 2)
        for umi1, umi2 in iter_umi_pairs:
            if hamming_distance(umi1, umi2) <= threshold:
                adj_list[umi1].append(umi2)
                adj_list[umi2].append(umi1)

        return adj_list

    def _get_adj_list_directional(self, umis, counts, threshold=1):
        ''' identify all umis within the hamming distance threshold
        and where the counts of the first umi is > (2 * second umi counts)-1'''

        adj_list = {umi: [] for umi in umis}
        if len(umis) > 25:
            umi_length = len(umis[0])
            substr_idx = build_substr_idx(umis, umi_length, threshold)
            iter_umi_pairs = iter_nearest_neighbours(umis, substr_idx)
        else:
            iter_umi_pairs = itertools.combinations(umis, 2)
        for umi1, umi2 in iter_umi_pairs:
            if hamming_distance(umi1, umi2) <= threshold:
                if counts[umi1] >= (counts[umi2]*2)-1:
                    adj_list[umi1].append(umi2)
                if counts[umi2] >= (counts[umi1]*2)-1:
                    adj_list[umi2].append(umi1)

        return adj_list

    def _get_adj_list_null(self, umis, counts, threshold):
        ''' for methods which don't use a adjacency dictionary'''
        return None

    # "get_connected_components" methods #

    def _get_connected_components_adjacency(self, umis, graph, counts):
        ''' find the connected UMIs within an adjacency dictionary'''

        # TS: TO DO: Work out why recursive function doesn't lead to same
        # final output. Then uncomment below

        # if len(graph) < 10000:
        #    self.search = breadth_first_search_recursive
        # else:
        #    self.search = breadth_first_search

        found = set()
        components = list()

        for node in sorted(graph, key=lambda x: counts[x], reverse=True):
            if node not in found:
                # component = self.search(node, graph)
                component = breadth_first_search(node, graph)
                found.update(component)
                components.append(component)
        return components

    def _get_connected_components_null(self, umis, adj_list, counts):
        ''' for methods which don't use a adjacency dictionary'''
        return umis

    # "group" methods #

    def _group_unique(self, clusters, adj_list, counts):
        ''' return groups for unique method'''
        if len(clusters) == 1:
            groups = [clusters]
        else:
            groups = [[x] for x in clusters]

        return groups

    def _group_directional(self, clusters, adj_list, counts):
        ''' return groups for directional method'''

        observed = set()
        groups = []
        for cluster in clusters:
            if len(cluster) == 1:
                groups.append(list(cluster))
                observed.update(cluster)
            else:
                cluster = sorted(cluster, key=lambda x: counts[x],
                                 reverse=True)
                # need to remove any node which has already been observed
                temp_cluster = []
                for node in cluster:
                    if node not in observed:
                        temp_cluster.append(node)
                        observed.add(node)
                groups.append(temp_cluster)

        return groups

    def _group_adjacency(self, clusters, adj_list, counts):
        ''' return groups for adjacency method'''

        groups = []

        for cluster in clusters:
            if len(cluster) == 1:
                groups.append(list(cluster))

            else:
                observed = set()

                lead_umis = self._get_best_min_account(cluster,
                                                       adj_list, counts)
                observed.update(lead_umis)

                for lead_umi in lead_umis:
                    connected_nodes = set(adj_list[lead_umi])
                    groups.append([lead_umi] +
                                  list(connected_nodes - observed))
                    observed.update(connected_nodes)

        return groups

    def _group_cluster(self, clusters, adj_list, counts):
        ''' return groups for cluster or directional methods'''

        groups = []
        for cluster in clusters:
            groups.append(sorted(cluster, key=lambda x: counts[x],
                                 reverse=True))

        return groups

    def _group_percentile(self, clusters, adj_list, counts):
        ''' Return "groups" for the the percentile method. Note
        that grouping isn't really compatible with the percentile
        method. This just returns the retained UMIs in a structure similar
        to other methods '''

        retained_umis = self._get_best_percentile(clusters, counts)
        groups = [[x] for x in retained_umis]

        return groups

    def __init__(self, cluster_method="directional"):
        ''' select the required class methods for the cluster_method'''

        self.max_umis_per_position = 0
        self.total_umis_per_position = 0
        self.positions = 0

        if cluster_method == "adjacency":
            self.get_adj_list = self._get_adj_list_adjacency
            self.get_connected_components = self._get_connected_components_adjacency
            self.get_groups = self._group_adjacency

        elif cluster_method == "directional":
            self.get_adj_list = self._get_adj_list_directional
            self.get_connected_components = self._get_connected_components_adjacency
            self.get_groups = self._group_directional

        elif cluster_method == "cluster":
            self.get_adj_list = self._get_adj_list_adjacency
            self.get_connected_components = self._get_connected_components_adjacency
            self.get_groups = self._group_cluster

        elif cluster_method == "percentile":
            self.get_adj_list = self._get_adj_list_null
            self.get_connected_components = self._get_connected_components_null
            # percentile method incompatible with defining UMI groups
            self.get_groups = self._group_percentile

        elif cluster_method == "unique":
            self.get_adj_list = self._get_adj_list_null
            self.get_connected_components = self._get_connected_components_null
            self.get_groups = self._group_unique

    def __call__(self, umis, counts, threshold):
        '''Counts is a directionary that maps UMIs to their counts'''

        umis = list(umis)

        self.positions += 1

        number_of_umis = len(umis)

        self.total_umis_per_position += number_of_umis

        if number_of_umis > self.max_umis_per_position:
            self.max_umis_per_position = number_of_umis

        len_umis = [len(x) for x in umis]

        assert max(len_umis) == min(len_umis), (
            "not all umis are the same length(!):  %d - %d" % (
                min(len_umis), max(len_umis)))

        adj_list = self.get_adj_list(umis, counts, threshold)
        clusters = self.get_connected_components(umis, adj_list, counts)
        final_umis = [list(x) for x in
                      self.get_groups(clusters, adj_list, counts)]

        return final_umis

class ReadDeduplicator:
    '''This is a wrapper for applying the UMI methods to bundles of BAM reads.
    It is currently a pretty transparent wrapper on UMIClusterer. Basically
    taking a read bundle, extracting the UMIs and Counts, running UMIClusterer
    and returning the results along with annotated reads'''

    def __init__(self, cluster_method="directional"):

        self.UMIClusterer = UMIClusterer(cluster_method=cluster_method)

    def __call__(self, bundle, threshold):
        '''Process the the bundled reads according to the method specified
        in the constructor. Return signature is:

        reads, final_umis, umi_counts, topologies, nodes

        reads:        predicted best reads for deduplicated position
        final_umis:   list of predicted parent UMIs
        umi_counts:   Sum of read counts for reads represented by the
                      corresponding UMI
        '''

        umis = bundle.keys()
        counts = {umi: bundle[umi]["count"] for umi in umis}

        clusters = self.UMIClusterer(umis, counts, threshold)

        final_umis = [cluster[0] for cluster in clusters]
        umi_counts = [sum(counts[umi] for umi in cluster)
                      for cluster in clusters]
        reads = [bundle[umi]["read"] for umi in final_umis]

        return (reads, final_umis, umi_counts)

class TwoPassPairWriter:
    '''This class makes a note of reads that need their pair outputting
    before outputting.  When the chromosome changes, the reads on that
    chromosome are read again, and any mates of reads already output
    are written and removed from the list of mates to output. When
    close is called, this is performed for the last chormosome, and
    then an algorithm identicate to pysam's mate() function is used to
    retrieve any remaining mates.

    This means that if close() is not called, at least as contigs
    worth of mates will be missing. '''

    def __init__(self, infile, outfile, tags=False):
        self.infile = infile
        self.outfile = outfile
        self.read1s = set()
        self.chrom = None

    def write(self, read, unique_id=None, umi=None, unmapped=False):
        '''Check if chromosome has changed since last time. If it has, scan
        for mates. Write the read to outfile and save the identity for paired
        end retrieval'''

        if unmapped or read.mate_is_unmapped:
            self.outfile.write(read)
            return

        if not self.chrom == read.reference_name:
            self.write_mates()
            self.chrom = read.reference_name

        key = read.query_name, read.next_reference_name, read.next_reference_start
        self.read1s.add(key)

        self.outfile.write(read)

    def write_mates(self):
        '''Scan the current chromosome for matches to any of the reads stored
        in the read1s buffer'''
        if self.chrom is not None:
            logging.debug("Dumping %i mates for contig %s" % (
                len(self.read1s), self.chrom))

        for read in self.infile.fetch(reference=self.chrom, multiple_iterators=True):
            if any((read.is_unmapped, read.mate_is_unmapped, read.is_read1)):
                continue

            key = read.query_name, read.reference_name, read.reference_start
            if key in self.read1s:
                self.outfile.write(read)
                self.read1s.remove(key)

        logging.debug("%i mates remaining" % len(self.read1s))

    def close(self):
        '''Write mates for remaining chromsome. Search for matches to any
        unmatched reads'''

        self.write_mates()
        logging.info("Searching for mates for %i unmatched alignments" %
               len(self.read1s))

        found = 0
        for read in self.infile.fetch(until_eof=True, multiple_iterators=True):

            if read.is_unmapped:
                continue

            key = read.query_name, read.reference_name, read.reference_start
            if key in self.read1s:
                self.outfile.write(read)
                self.read1s.remove(key)
                found += 1
                continue

        logging.info("%i mates never found" % len(self.read1s))
        self.outfile.close()
    
class get_bundles:

    ''' A functor - When called returns a dictionary of dictionaries,
    representing the unique reads at a position/spliced/strand
    combination. The key to the top level dictionary is a umi. Each
    dictionary contains a "read" entry with the best read, and a count
    entry with the number of reads with that
    position/spliced/strand/umi combination

    initiation arguments:

    options: script options

    all_reads: if true, return all reads in the dictionary. Else,
    return the 'best' read (using MAPQ +/- multimapping) for each key

    return_read2: Return read2s immediately as a single read

    metacontig_contig: Maps metacontigs to the consistuent contigs
    '''

    def __init__(self,
                 options,
                 only_count_reads=False,
                 all_reads=False,
                 return_unmapped=False,
                 return_read2=False,
                 metacontig_contig=None):

        self.options = options
        self.only_count_reads = only_count_reads
        self.all_reads = all_reads
        self.return_unmapped = return_unmapped
        self.return_read2 = return_read2
        self.metacontig_contig = metacontig_contig

        self.contig_metacontig = {}
        if self.metacontig_contig:
            for metacontig in metacontig_contig:
                for contig in metacontig_contig[metacontig]:
                    self.contig_metacontig[contig] = metacontig

        # set the method with which to extract umis from reads
        if self.options.get_umi_method == "read_id":
            self.barcode_getter = partial(
                get_barcode_read_id,
                cell_barcode=self.options.per_cell,
                sep=self.options.umi_sep)
        else:
            raise ValueError("Unknown UMI extraction method")

        self.read_events = collections.Counter()
        self.observed_contigs = collections.defaultdict(set)

        self.last_pos = 0
        self.last_chr = None
        self.start = 0
        self.current_chr = None
        self.last_umiStart=0

        self.reads_dict = collections.defaultdict(
            lambda: collections.defaultdict(
                lambda: collections.defaultdict(dict)))
        self.read_counts = collections.defaultdict(
            lambda: collections.defaultdict(dict))

    def update_dicts(self, read, pos, key, umi):

        # The content of the reads_dict depends on whether all reads
        # are being retained

        if self.all_reads:
            # retain all reads per key
            try:
                self.reads_dict[pos][key][umi]["count"] += 1
            except KeyError:
                self.reads_dict[pos][key][umi]["read"] = [read]
                self.reads_dict[pos][key][umi]["count"] = 1
            else:
                self.reads_dict[pos][key][umi]["read"].append(read)

        elif self.only_count_reads:
            # retain all reads per key
            try:
                self.reads_dict[pos][key][umi]["count"] += 1
            except KeyError:
                self.reads_dict[pos][key][umi]["count"] = 1

        else:
            # retain just a single read per key
            try:
                self.reads_dict[pos][key][umi]["count"] += 1
            except KeyError:
                self.reads_dict[pos][key][umi]["read"] = read
                self.reads_dict[pos][key][umi]["count"] = 1
                self.read_counts[pos][key][umi] = 0
            else:
                old_read=self.reads_dict[pos][key][umi]["read"]
                if read.get_tag("NM")<=old_read.get_tag("NM"):
                    self.reads_dict[pos][key][umi]["read"] = read
                    self.read_counts[pos][key][umi] = 0
                    return

                # TS: implemented different checks for multimapping here
                if self.options.detection_method in ["NH", "X0"]:
                    tag = self.options.detection_method
                    if (self.reads_dict[pos][key][umi]["read"].opt(tag) <
                        read.opt(tag)):
                        return
                    elif (self.reads_dict[pos][key][umi]["read"].opt(tag) >
                          read.opt(tag)):
                        self.reads_dict[pos][key][umi]["read"] = read
                        self.read_counts[pos][key][umi] = 0

                elif self.options.detection_method == "XT":
                    if self.reads_dict[pos][key][umi]["read"].opt("XT") == "U":
                        return
                    elif read.opt("XT") == "U":
                        self.reads_dict[pos][key][umi]["read"] = read
                        self.read_counts[pos][key][umi] = 0

                self.read_counts[pos][key][umi] += 1
                prob = 1.0/self.read_counts[pos][key][umi]

                #if random.random() < prob:
                #    self.reads_dict[pos][key][umi]["read"] = read

    def check_output(self):

        do_output = False
        out_keys = None

        if self.options.per_gene:

            if self.metacontig_contig:

                if (self.current_chr != self.last_chr and
                    (self.observed_contigs[self.last_pos] ==
                     self.metacontig_contig[self.last_pos])):
                    do_output = True
                    out_keys = [self.last_pos]

            else:
                if self.current_chr != self.last_chr:
                    do_output = True
                    out_keys = sorted(self.reads_dict.keys())

        elif self.options.whole_contig:

            if self.current_chr != self.last_chr:
                do_output = True
                out_keys = sorted(self.reads_dict.keys())

        else:

            if (self.start > (self.last_pos+1000) or
                self.current_chr != self.last_chr):
                self.shift_umis(1)
                do_output = True
                out_keys = sorted(self.reads_dict.keys())
                
                if self.current_chr == self.last_chr:
                    out_keys = [x for x in out_keys if x <= self.start-1000]

        return do_output, out_keys

    def compare_umis(self,p,k,p_near,k_near):
        umi=self.reads_dict[p][k].keys()
        umi_near=self.reads_dict[p_near][k_near].keys()
        umi_overlap = set(umi).intersection(set(umi_near))
        if len(umi_overlap)>0:
            for u in umi_overlap:
                near_count=self.reads_dict[p_near][k_near][u]["count"]
                p_count=self.reads_dict[p][k][u]["count"]
                near_read=self.reads_dict[p_near][k_near][u]["read"]
                p_read=self.reads_dict[p][k][u]["read"]
                if near_count > p_count:
                    self.reads_dict[p_near][k_near][u]["count"]+=p_count
                    del self.reads_dict[p][k][u]
                elif near_count < p_count:
                    self.reads_dict[p][k][u]["count"]+=near_count
                    del self.reads_dict[p_near][k_near][u]
                else:
                    if p_read.get_tag("NM") < near_read.get_tag("NM"):
                        self.reads_dict[p][k][u]["count"]+=near_count
                        del self.reads_dict[p_near][k_near][u]
                    elif p_read.get_tag("NM") > near_read.get_tag("NM"):
                        self.reads_dict[p_near][k_near][u]["count"]+=p_count
                        del self.reads_dict[p][k][u]
                    else:
                        if k[0][0]:
                            self.reads_dict[p_near][k_near][u]["count"]+=p_count
                            del self.reads_dict[p][k][u]
                        else:
                            self.reads_dict[p][k][u]["count"]+=near_count
                            del self.reads_dict[p_near][k_near][u]

    def shift_umis(self, shift_bp):
        for p in sorted(self.reads_dict.keys()):
            #umi_collector=lambda: collections.defaultdict(dict)
            p_near=p+shift_bp
            p_keys=self.reads_dict[p].keys()
            p_keys.sort(key=lambda x:x[0][2])
            #print p_keys
            for k in p_keys:
                if self.options.paired == False:
                    if p_near in self.reads_dict.keys():
                        self.compare_umis(p,k,p_near,k)
                else:
                    k_near=((k[0][0],k[0][1],k[0][2]+shift_bp,k[0][3]),None)
                    if k_near in self.reads_dict[p].keys():
                        self.compare_umis(p,k_near,p,k)
                    if p_near in self.reads_dict.keys():
                        k_near=((k[0][0],k[0][1],k[0][2]-shift_bp,k[0][3]),None)
                        if k_near in self.reads_dict[p_near].keys():
                            self.compare_umis(p,k,p_near,k_near)
                        if k in self.reads_dict[p_near].keys():
                            self.compare_umis(p,k,p_near,k)


    def __call__(self, inreads):

        for read in inreads:

            if read.is_read2:
                if self.return_read2:
                    if not read.is_unmapped or (
                            read.is_unmapped and self.return_unmapped):
                        yield read, None, "single_read"
                continue
            else:
                self.read_events['Input Reads'] += 1

            if read.is_unmapped:
                if self.options.paired:
                    if read.mate_is_unmapped:
                        self.read_events['Both unmapped'] += 1
                    else:
                        self.read_events['Read 1 unmapped'] += 1
                else:
                    self.read_events['Single end unmapped'] += 1

                if self.return_unmapped:
                    self.read_events['Input Reads'] += 1
                    yield read, None, "single_read"
                continue

            if read.mate_is_unmapped and self.options.paired:
                if not read.is_unmapped:
                    self.read_events['Read 2 unmapped'] += 1
                if self.return_unmapped:
                    yield read, None, "single_read"
                continue

            if self.options.paired:
                self.read_events['Paired Reads'] += 1

            if self.options.subset:
                if random.random() >= self.options.subset:
                    self.read_events['Randomly excluded'] += 1
                    continue

            if self.options.mapping_quality:
                if read.mapq < self.options.mapping_quality:
                    self.read_events['< MAPQ threshold'] += 1
                    continue

            self.current_chr = read.reference_name

            if self.options.per_gene:

                if self.options.per_contig:

                    if self.metacontig_contig:
                        transcript = read.reference_name
                        gene = self.contig_metacontig[transcript]
                    else:
                        gene = read.reference_name

                elif self.options.gene_tag:

                    try:
                        gene = read.get_tag(self.options.gene_tag)
                    except KeyError:
                        self.read_events['Read skipped, no tag'] += 1
                        continue

                    if re.search(self.options.skip_regex, gene):
                        self.read_events['Gene skipped - matches regex'] += 1
                        continue

                pos = gene
                key = pos

                if self.last_chr:
                    do_output, out_keys = self.check_output()
                else:
                    do_output = False

                if do_output:
                    for p in out_keys:
                        for k in sorted(self.reads_dict[p].keys()):
                            yield self.reads_dict[p][k], k, "bundle"

                        del self.reads_dict[p]

                self.last_chr = self.current_chr
                self.last_pos = pos

            else:
                start, pos, is_spliced = get_read_position(
                    read, self.options.soft_clip_threshold)

                do_output, out_keys = self.check_output()

                if do_output:
                    for p in out_keys:
                        for k in sorted(self.reads_dict[p].keys()):
                            if len(self.reads_dict[p][k].keys())>0:
                                yield self.reads_dict[p][k], k, "bundle"

                        del self.reads_dict[p]
                        if p in self.read_counts:
                            del self.read_counts[p]

                self.last_pos = self.start
                self.last_chr = self.current_chr
                
                if self.options.read_length:
                    r_length = read.query_length
                else:
                    r_length = 0
                    
                key = (read.is_reverse, self.options.spliced & is_spliced,
                       self.options.paired*read.tlen, r_length)

            # get the umi +/- cell barcode and update dictionaries
            if self.options.ignore_umi:
                if self.options.per_cell:
                    umi, cell = self.barcode_getter(read)
                    umi = ""
                else:
                    umi, cell = "", ""
            else:
                umi, cell = self.barcode_getter(read)
            
            key = (key, cell)
            self.update_dicts(read, pos, key, umi)

            if self.metacontig_contig:
                # keep track of observed contigs for each gene
                self.observed_contigs[gene].add(transcript)

        # yield remaining bundles
        self.shift_umis(1)
        for p in sorted(self.reads_dict.keys()):
            for k in sorted(self.reads_dict[p].keys()):
                if len(self.reads_dict[p][k].keys())>0:
                    yield self.reads_dict[p][k], k, "bundle"

class BetterFormatter(optparse.IndentedHelpFormatter):
    """A formatter for :class:`OptionParser` outputting indented
    help text.
    """

    def __init__(self, *args, **kwargs):

        optparse.IndentedHelpFormatter.__init__(self, *args, **kwargs)
        self.wrapper = textwrap.TextWrapper(width=self.width)

    def _formatter(self, text):

        return '\n'.join(['\n'.join(p) for p in
                          map(self.wrapper.wrap,
                              self.parser.expand_prog_name(text).split('\n'))])

    def format_description(self, description):

        if description:
            return self._formatter(description) + '\n'
        else:
            return ''

    def format_epilog(self, epilog):

        if epilog:
            return '\n' + self._formatter(epilog) + '\n'
        else:
            return ''

    def format_usage(self, usage):

        return self._formatter(optparse._("Usage: %s\n") % usage)

    def format_option(self, option):

        # Ripped and modified from Python 2.6's optparse's HelpFormatter
        result = []
        opts = self.option_strings[option]
        opt_width = self.help_position - self.current_indent - 2
        if len(opts) > opt_width:
            opts = "%*s%s\n" % (self.current_indent, "", opts)
            indent_first = self.help_position
        else:                       # start help on same line as opts
            opts = "%*s%-*s  " % (self.current_indent, "", opt_width, opts)
            indent_first = 0
        result.append(opts)

        if option.help:
            help_text = self.expand_default(option)
            # Added expand program name
            help_text = self.parser.expand_prog_name(help_text)
            # Modified the generation of help_line
            help_lines = []
            wrapper = textwrap.TextWrapper(width=self.help_width)
            for p in map(wrapper.wrap, help_text.split('\n')):
                if p:
                    help_lines.extend(p)
                else:
                    help_lines.append('')
            # End of modification
            result.append("%*s%s\n" % (indent_first, "", help_lines[0]))
            result.extend(["%*s%s\n" % (self.help_position, "", line)
                           for line in help_lines[1:]])
        elif opts[-1] != "\n":
            result.append("\n")

        return "".join(result)

class AppendCommaOption(optparse.Option):
    def convert_value(self, opt, value):
        if value is not None:
            if self.nargs == 1:
                if self.action == "append":
                    if "," in value:
                        return [self.check_value(opt, v) for v in
                                value.split(",") if v != ""]
                    else:
                        if value != "":
                            return self.check_value(opt, value)
                        else:
                            return value
                else:
                    return self.check_value(opt, value)
            else:
                return tuple([self.check_value(opt, v) for v in value])

    # why is it necessary to pass action and dest to this function when
    # they could be accessed as self.action and self.dest?
    def take_action(self, action, dest, opt, value, values, parser):

        if action == "append" and type(value) == list:
            values.ensure_value(dest, []).extend(value)
        else:
            optparse.Option.take_action(
                self, action, dest, opt, value, values, parser)

class OptionParser(optparse.OptionParser):

    '''UMI-tools derivative of OptionParser.
    '''

    def __init__(self, *args, **kwargs):
        # if "--short" is a command line option
        # remove usage from kwargs
        if "--no-usage" in sys.argv:
            kwargs["usage"] = None

        optparse.OptionParser.__init__(self, *args,
                                       option_class=AppendCommaOption,
                                       formatter=BetterFormatter(),
                                       **kwargs)

        # set new option parser
        # parser.formatter = BetterFormatter()
        # parser.formatter.set_parser(parser)
        if "--no-usage" in sys.argv:
            self.add_option("--no-usage", dest="help_no_usage",
                            action="store_true",
                            help="output help without usage information")

class OptionGroup(optparse.OptionGroup):
    pass

class MultiLineFormatter(logging.Formatter):

    '''logfile formatter: add identation for multi-line entries.'''

def callbackShortHelp(option, opt, value, parser):
    '''output short help (only command line options).'''
    # clear usage and description
    parser.set_description(None)
    parser.set_usage(None)
    # output help
    parser.print_help()
    # exit
    parser.exit()

def openFile(filename, mode="r", create_dir=False):
    '''open file in *filename* with mode *mode*.

    If *create* is set, the directory containing filename
    will be created if it does not exist.

    gzip - compressed files are recognized by the
    suffix ``.gz`` and opened transparently.

    Note that there are differences in the file
    like objects returned, for example in the
    ability to seek.

    returns a file or file-like object.
    '''

    _, ext = os.path.splitext(filename)

    if create_dir:
        dirname = os.path.dirname(filename)
        if dirname and not os.path.exists(dirname):
            os.makedirs(dirname)

    if ext.lower() in (".gz", ".z"):
        if sys.version_info.major >= 3:
            if mode == "r":
                return gzip.open(filename, 'rt', encoding="ascii")
            elif mode == "w":
                return gzip.open(filename, 'wt',
                                 compresslevel=global_options.compresslevel,
                                 encoding="ascii")
            else:
                raise NotImplementedError(
                    "mode '{}' not implemented".format(mode))
        else:
            return gzip.open(filename, mode,
                             compresslevel=global_options.compresslevel)
    else:
        return open(filename, mode)

def getHeader():
    """return a header string with command line options and timestamp

    """
    system, host, release, version, machine = os.uname()

    return "# output generated by %s\n# job started at %s on %s -- %s\n# pid: %i, system: %s %s %s %s" %\
           (" ".join(sys.argv),
            time.asctime(time.localtime(time.time())),
            host,
            global_id,
            os.getpid(),
            system, release, version, machine)

def getParams(options=None):
    """return a string containing script parameters.

    Parameters are all variables that start with ``param_``.
    """
    result = []
    if options:
        members = options.__dict__
        for k, v in sorted(members.items()):
            result.append("# %-40s: %s" % (k, str(v)))
    else:
        vars = inspect.currentframe().f_back.f_locals
        for var in filter(lambda x: re.match("param_", x), vars.keys()):
            result.append("# %-40s: %s" %
                          (var, str(vars[var])))

    if result:
        return "\n".join(result)
    else:
        return "# no parameters."

def getFooter():
    """return a header string with command line options and
    timestamp.
    """
    return "# job finished in %i seconds at %s -- %s -- %s" %\
           (time.time() - global_starting_time,
            time.asctime(time.localtime(time.time())),
            " ".join(map(lambda x: "%5.2f" % x, os.times()[:4])),
            global_id)


def Start(parser=None,
          argv=sys.argv,
          quiet=False,
          add_pipe_options=True,
          add_group_dedup_options=True,
          add_sam_options=True,
          return_parser=False):
    if not parser:
        parser = OptionParser(
            version="%prog version: $Id$")

    global global_options, global_args, global_starting_time

    # save default values given by user
    user_defaults = copy.copy(parser.defaults)

    global_starting_time = time.time()

    if add_sam_options:
        group = OptionGroup(parser, "Input options")

        group.add_option("-i", "--in-sam", dest="in_sam", action="store_true",
                         help="Input file is in sam format [default=%default]",
                         default=False)

        group.add_option("--extract-umi-method", dest="get_umi_method", type="choice",
                         choices=("read_id", "tag", "umis"), default="read_id",
                         help="how is the read UMI +/ cell barcode encoded? "
                         "[default=%default]")

        group.add_option("--umi-separator", dest="umi_sep",
                         type="string", help="separator between read id and UMI",
                         default="_")

        group.add_option("--umi-tag", dest="umi_tag",
                         type="string", help="tag containing umi",
                         default='RX')

        group.add_option("--umi-tag-split", dest="umi_tag_split",
                         type="string",
                         help="split UMI in tag and take the first element",
                         default=None)

        group.add_option("--umi-tag-delimiter", dest="umi_tag_delim",
                         type="string",
                         help="concatenate UMI in tag separated by delimiter",
                         default=None)

        group.add_option("--cell-tag", dest="cell_tag",
                         type="string", help="tag containing cell barcode",
                         default=None)

        group.add_option("--cell-tag-split", dest="cell_tag_split",
                         type="string",
                         help=("split cell barcode in tag and take the first element"
                               "for e.g 10X GEM tags"),
                         default='-')

        group.add_option("--cell-tag-delimiter", dest="cell_tag_delim",
                         type="string",
                         help="concatenate cell barcode in tag separated by delimiter",
                         default=None)

        group.add_option("--paired", dest="paired", action="store_true",
                         default=False,
                         help="paired BAM. [default=%default]")

        group.add_option("--mapping-quality", dest="mapping_quality",
                         type="int",
                         help="Minimum mapping quality for a read to be retained"
                         " [default=%default]",
                         default=0)

        parser.add_option_group(group)

        group = OptionGroup(parser, "UMI grouping options")

        group.add_option("--method", dest="method", type="choice",
                         choices=("adjacency", "directional",
                                  "percentile", "unique", "cluster"),
                         default="directional",
                         help="method to use for umi grouping [default=%default]")

        group.add_option("--edit-distance-threshold", dest="threshold",
                         type="int",
                         default=1,
                         help="Edit distance theshold at which to join two UMIs "
                         "when grouping UMIs. [default=%default]")

        parser.add_option_group(group)

        group = OptionGroup(parser, "Single-cell RNA-Seq options")

        group.add_option("--per-gene", dest="per_gene", action="store_true",
                         default=False,
                         help="Group/Dedup/Count per gene. Must combine with "
                         "either --gene-tag or --per-contig")

        group.add_option("--gene-tag", dest="gene_tag",
                         type="string",
                         help="gene is defined by this bam tag [default=%default]",
                         default=None)

        group.add_option("--skip-tags-regex", dest="skip_regex",
                         type="string",
                         help="Used with --gene-tag. "
                         "Ignore reads where the gene-tag matches this regex",
                         default="^(__|Unassigned)")

        group.add_option("--per-contig", dest="per_contig", action="store_true",
                         default=False,
                         help="count per contig (field 3 in BAM; RNAME),"
                         " e.g for transcriptome where contig = gene")

        group.add_option("--gene-transcript-map", dest="gene_transcript_map",
                         type="string",
                         help="file mapping transcripts to genes (tab separated)",
                         default=None)

        group.add_option("--per-cell", dest="per_cell", action="store_true",
                         default=False,
                         help="Group/Dedup/Count per cell")

        parser.add_option_group(group)

    if add_group_dedup_options:

        group = OptionGroup(parser, "Group/Dedup options")

        group.add_option("-o", "--out-sam", dest="out_sam", action="store_true",
                         help="Output alignments in sam format [default=%default]",
                         default=False)

        group.add_option("--no-sort-output", dest="no_sort_output",
                         action="store_true", default=False,
                         help="Don't Sort the output")

        group.add_option("--buffer-whole-contig", dest="whole_contig",
                         action="store_true", default=False,
                         help="Read whole contig before outputting bundles: "
                         "guarantees that no reads are missed, but increases "
                         "memory usage")

        group.add_option("--whole-contig", dest="whole_contig",
                         action="store_true", default=False,
                         help=optparse.SUPPRESS_HELP)

        group.add_option("--multimapping-detection-method",
                         dest="detection_method", type="choice",
                         choices=("NH", "X0", "XT"),
                         default=None,
                         help="Some aligners identify multimapping using bam "
                         "tags. Setting this option to NH, X0 or XT will "
                         "use these tags when selecting the best read "
                         "amongst reads with the same position and umi "
                         "[default=%default]")

        group.add_option("--spliced-is-unique", dest="spliced",
                         action="store_true",
                         help="Treat a spliced read as different to an unspliced"
                         " one [default=%default]",
                         default=False)

        group.add_option("--soft-clip-threshold", dest="soft_clip_threshold",
                         type="float",
                         help="number of bases clipped from 5' end before"
                         "read is counted as spliced [default=%default]",
                         default=4)

        group.add_option("--read-length", dest="read_length",
                         action="store_true", default=False,
                         help="use read length in addition to position and UMI"
                         "to identify possible duplicates [default=%default]")

        parser.add_option_group(group)

    # options added separately here to maintain better output order
    if add_sam_options:
        group = OptionGroup(parser, "debug options")

        group.add_option("--ignore-umi", dest="ignore_umi",
                         action="store_true", help="Ignore UMI and dedup"
                         " only on position", default=False)

        group.add_option("--chrom", dest="chrom", type="string",
                         help="Restrict to one chromosome",
                         default=None)

        group.add_option("--subset", dest="subset", type="float",
                         help="Use only a fraction of reads, specified by subset",
                         default=None)

        parser.add_option_group(group)

    group = OptionGroup(parser, "profiling options")

    group.add_option("--timeit", dest='timeit_file', type="string",
                     help="store timeing information in file [%default].")

    group.add_option("--timeit-name", dest='timeit_name', type="string",
                     help="name in timing file for this class of jobs "
                     "[%default].")

    group.add_option("--timeit-header", dest='timeit_header',
                     action="store_true",
                     help="add header for timing information [%default].")

    parser.add_option_group(group)

    group = OptionGroup(parser, "common options")

    group.add_option("-v", "--verbose", dest="loglevel", type="int",
                     help="loglevel [%default]. The higher, the more output.")

    group.add_option("-?", dest="short_help", action="callback",
                     callback=callbackShortHelp,
                     help="output short help (command line options only.")

    group.add_option("--random-seed", dest='random_seed', type="int",
                     help="random seed to initialize number generator "
                     "with [%default].")

    parser.add_option_group(group)

    if quiet:
        parser.set_defaults(loglevel=0)
    else:
        parser.set_defaults(loglevel=1)

    parser.set_defaults(
        timeit_file=None,
        timeit_name='all',
        timeit_header=None,
        random_seed=None,
    )

    if add_pipe_options:
        group = OptionGroup(parser, "Input/output options")
        group.add_option("-I", "--stdin", dest="stdin", type="string",
                         help="file to read stdin from [default = stdin].",
                         metavar="FILE")
        group.add_option("-L", "--log", dest="stdlog", type="string",
                         help="file with logging information "
                         "[default = stdout].",
                         metavar="FILE")
        group.add_option("-E", "--error", dest="stderr", type="string",
                         help="file with error information "
                         "[default = stderr].",
                         metavar="FILE")
        group.add_option("-S", "--stdout", dest="stdout", type="string",
                         help="file where output is to go "
                         "[default = stdout].",
                         metavar="FILE")
        group.add_option("--log2stderr", dest="log2stderr",
                         action="store_true", help="send logging information"
                         " to stderr [default = False].")
        group.add_option("--compresslevel", dest="compresslevel", type="int",
                         help="Level of Gzip compression to use. Default (6) matches"
                         "GNU gzip rather than python gzip default (which is 9)")

        parser.set_defaults(stderr=sys.stderr)
        parser.set_defaults(stdout=sys.stdout)
        parser.set_defaults(stdlog=sys.stdout)
        parser.set_defaults(stdin=sys.stdin)
        parser.set_defaults(log2stderr=False)
        parser.set_defaults(compresslevel=6)

    parser.add_option_group(group)

    # restore user defaults
    parser.defaults.update(user_defaults)

    if return_parser:
        return parser

    global_options, global_args = parser.parse_args(argv[1:])

    if global_options.random_seed is not None:
        random.seed(global_options.random_seed)

    if add_pipe_options:
        if global_options.stdout != sys.stdout:
            global_options.stdout = openFile(global_options.stdout, "w")
        if global_options.stderr != sys.stderr:
            if global_options.stderr == "stderr":
                global_options.stderr = global_options.stderr
            else:
                global_options.stderr = openFile(global_options.stderr, "w")
        if global_options.stdlog != sys.stdout:
            global_options.stdlog = openFile(global_options.stdlog, "a")
        elif global_options.log2stderr:
            global_options.stdlog = global_options.stderr
        if global_options.stdin != sys.stdin:
            global_options.stdin = openFile(global_options.stdin, "r")
    else:
        global_options.stderr = sys.stderr
        global_options.stdout = sys.stdout
        global_options.stdin = sys.stdin
        if global_options.log2stderr:
            global_options.stdlog = sys.stderr
        else:
            global_options.stdlog = sys.stdout

    if global_options.loglevel >= 1:
        global_options.stdlog.write(getHeader() + "\n")
        global_options.stdlog.write(getParams(global_options) + "\n")
        global_options.stdlog.flush()

    # configure logging
    # map from 0-10 to logging scale
    # 0: quiet
    # 1: little verbositiy
    # >1: increased verbosity
    if global_options.loglevel == 0:
        lvl = logging.ERROR
    elif global_options.loglevel == 1:
        lvl = logging.INFO
    else:
        lvl = logging.DEBUG

    if global_options.stdout == global_options.stdlog:
        format = '# %(asctime)s %(levelname)s %(message)s'
    else:
        format = '%(asctime)s %(levelname)s %(message)s'

    logging.basicConfig(
        level=lvl,
        format=format,
        stream=global_options.stdlog)

    # set up multi-line logging
    # Note that .handlers is not part of the API, might change
    # Solution is to configure handlers explicitely.
    for handler in logging.getLogger().handlers:
        handler.setFormatter(MultiLineFormatter(format))

    return global_options, global_args

def validateSamOptions(options):
    ''' Check the validity of the option combinations '''

    if options.per_gene:
        if options.gene_tag and options.per_contig:
            raise ValueError("need to use either --per-contig "
                             "OR --gene-tag, please do not provide both")

        if not options.per_contig and not options.gene_tag:
            raise ValueError("for per-gene applications, must supply "
                             "--per-contig or --gene-tag")

    if options.per_contig and not options.per_gene:
        raise ValueError("need to use --per-gene with --per-contig")

    if options.gene_tag and not options.per_gene:
        raise ValueError("need to use --per-gene with --gene_tag")

    if options.gene_transcript_map and not options.per_contig:
        raise ValueError("need to use --per-contig and --per-gene"
                         "with --gene-transcript-map")

    if options.get_umi_method == "tag":
        if options.umi_tag is None:
            raise ValueError("Need to supply the --umi-tag option")
        if options.per_cell and options.cell_tag is None:
            raise ValueError("Need to supply the --cell-tag option")

    if options.skip_regex:
        try:
            re.compile(options.skip_regex)
        except re.error:
            raise ValueError("skip-regex '%s' is not a "
                             "valid regex" % options.skip_regex)

def Stop():
    """stop the experiment.

    This method performs final book-keeping, closes the output streams
    and writes the final log messages indicating script completion.
    """

    if global_options.loglevel >= 1 and global_benchmark:
        t = time.time() - global_starting_time
        global_options.stdlog.write(
            "######### Time spent in benchmarked functions #########\n")
        global_options.stdlog.write("# function\tseconds\tpercent\n")
        for key, value in global_benchmark.items():
            global_options.stdlog.write(
                "# %s\t%6i\t%5.2f%%\n" % (key, value,
                                          (100.0 * float(value) / t)))
        global_options.stdlog.write(
            "#######################################################\n")

    if global_options.loglevel >= 1:
        global_options.stdlog.write(getFooter() + "\n")

    # close files
    if global_options.stdout != sys.stdout:
        global_options.stdout.close()
    # do not close log, otherwise error occurs in atext.py
    # if global_options.stdlog != sys.stdout:
    #   global_options.stdlog.close()

    if global_options.stderr != sys.stderr:
        global_options.stderr.close()

    if global_options.timeit_file:

        outfile = open(global_options.timeit_file, "a")

        if global_options.timeit_header:
            outfile.write("\t".join(
                ("name", "wall", "user", "sys", "cuser", "csys",
                 "host", "system", "release", "machine",
                 "start", "end", "path", "cmd")) + "\n")

        csystem, host, release, version, machine = map(str, os.uname())
        uusr, usys, c_usr, c_sys = map(lambda x: "%5.2f" % x, os.times()[:4])
        t_end = time.time()
        c_wall = "%5.2f" % (t_end - global_starting_time)

        if sys.argv[0] == "run.py":
            cmd = global_args[0]
            if len(global_args) > 1:
                cmd += " '" + "' '".join(global_args[1:]) + "'"
        else:
            cmd = sys.argv[0]

        result = "\t".join((global_options.timeit_name,
                            c_wall, uusr, usys, c_usr, c_sys,
                            host, csystem, release, machine,
                            time.asctime(time.localtime(global_starting_time)),
                            time.asctime(time.localtime(t_end)),
                            os.path.abspath(os.getcwd()),
                            cmd)) + "\n"

        outfile.write(result)
        outfile.close()

def info(message):
    '''log information message, see the :mod:`logging` module'''
    logging.info(message)

def warn(message):
    '''log warning message, see the :mod:`logging` module'''
    logging.warning(message)

def getTempFile(dir=None, shared=False, suffix=""):
    return tempfile.NamedTemporaryFile(dir=dir, delete=False, prefix="ctmp",suffix=suffix)

def getTempFilename(dir=None, shared=False, suffix=""):
    tmpfile = getTempFile(dir=dir, shared=shared, suffix=suffix)
    tmpfile.close()
    return tmpfile.name

def main(argv=None):
    """script main.

    parses command line options in sys.argv, unless *argv* is given.
    """

    if argv is None:
        argv = sys.argv

    # setup command line parser
    parser = OptionParser(version="%prog version: $Id$",
                            usage=globals()["__doc__"])
    group = OptionGroup(parser, "dedup-specific options")

    group.add_option("--output-stats", dest="stats", type="string",
                     default=False,
                     help="Specify location to output stats")

    parser.add_option_group(group)

    # add common options (-h/--help, ...) and parse command line
    (options, args) = Start(parser, argv=argv)

    validateSamOptions(options)

    if options.random_seed:
        np.random.seed(options.random_seed)

    if options.stdin != sys.stdin:
        in_name = options.stdin.name
        options.stdin.close()
    else:
        raise ValueError("Input on standard in not currently supported")

    if options.stdout != sys.stdout:
        if options.no_sort_output:
            out_name = options.stdout.name
        else:
            out_name = getTempFilename()
            sorted_out_name = options.stdout.name
        options.stdout.close()
    else:
        if options.no_sort_output:
            out_name = "-"
        else:
            out_name = getTempFilename()
            sorted_out_name = "-"

    if not options.no_sort_output:  # need to determine the output format for sort
        if options.out_sam:
            sort_format = "sam"
        else:
            sort_format = "bam"

    if options.in_sam:
        in_mode = "r"
    else:
        in_mode = "rb"

    if options.out_sam:
        out_mode = "wh"
    else:
        out_mode = "wb"

    if options.stats and options.ignore_umi:
        raise ValueError("'--output-stats' and '--ignore-umi' options"
                         " cannot be used together")

    infile = pysam.Samfile(in_name, in_mode)
    outfile = pysam.Samfile(out_name, out_mode, template=infile)

    if options.paired:
        outfile = TwoPassPairWriter(infile, outfile)

    nInput, nOutput, input_reads, output_reads = 0, 0, 0, 0
    
    gene_tag = options.gene_tag
    metacontig2contig = None

    if options.chrom:
        inreads = infile.fetch(reference=options.chrom)

    else:
        if options.per_contig and options.gene_transcript_map:
            '''
            metacontig2contig = getMetaContig2contig(
                infile, options.gene_transcript_map)
            metatag = "MC"
            inreads = metafetcher(infile, metacontig2contig, metatag)
            gene_tag = metatag
            '''

        else:
            inreads = infile.fetch()

    # set up ReadCluster functor with methods specific to
    # specified options.method
    processor = ReadDeduplicator(options.method)

    bundle_iterator = get_bundles(
        options,
        metacontig_contig=metacontig2contig)
    '''
    if options.stats:
        # set up arrays to hold stats data
        stats_pre_df_dict = {"UMI": [], "counts": []}
        stats_post_df_dict = {"UMI": [], "counts": []}
        pre_cluster_stats = []
        post_cluster_stats = []
        pre_cluster_stats_null = []
        post_cluster_stats_null = []
        topology_counts = collections.Counter()
        node_counts = collections.Counter()
        read_gn = random_read_generator(
            infile.filename, chrom=options.chrom,
            barcode_getter=bundle_iterator.barcode_getter)
    '''
    for bundle, key, status in bundle_iterator(inreads):

        nInput += sum([bundle[umi]["count"] for umi in bundle])

        while nOutput >= output_reads + 100000:
            output_reads += 100000
            info("Written out %i reads" % output_reads)

        while nInput >= input_reads + 1000000:
            input_reads += 1000000
            info("Parsed %i input reads" % input_reads)
        '''
        if options.stats:
            # generate pre-dudep stats
            average_distance = get_average_umi_distance(bundle.keys())
            pre_cluster_stats.append(average_distance)
            cluster_size = len(bundle)
            random_umis = read_gn.getUmis(cluster_size)
            average_distance_null = get_average_umi_distance(random_umis)
            pre_cluster_stats_null.append(average_distance_null)
        '''
        if options.ignore_umi:
            for umi in bundle:
                nOutput += 1
                outfile.write(bundle[umi]["read"])

        else:

            # dedup using umis and write out deduped bam
            reads, umis, umi_counts = processor(
                bundle=bundle,
                threshold=options.threshold)

            for read in reads:
                outfile.write(read)
                nOutput += 1
            '''
            if options.stats:

                # collect pre-dudupe stats
                stats_pre_df_dict['UMI'].extend(bundle)
                stats_pre_df_dict['counts'].extend(
                    [bundle[UMI]['count'] for UMI in bundle])

                # collect post-dudupe stats
                post_cluster_umis = [bundle_iterator.barcode_getter(x)[0] for x in reads]
                stats_post_df_dict['UMI'].extend(umis)
                stats_post_df_dict['counts'].extend(umi_counts)

#                average_distance = get_average_umi_distance(post_cluster_umis)
                post_cluster_stats.append(average_distance)

                cluster_size = len(post_cluster_umis)
                random_umis = read_gn.getUmis(cluster_size)
                average_distance_null = get_average_umi_distance(random_umis)
                post_cluster_stats_null.append(average_distance_null)
            '''
    outfile.close()

    if not options.no_sort_output:
        # sort the output
        pysam.sort("-o", sorted_out_name, "-O", sort_format, out_name)
        os.unlink(out_name)  # delete the tempfile

    # write footer and output benchmark information.
    info(
        "Reads: %s" % ", ".join(["%s: %s" % (x[0], x[1]) for x in
                                 bundle_iterator.read_events.most_common()]))

    info("Number of reads out: %i" % nOutput)

    if not options.ignore_umi:  # otherwise processor has not been used
        info("Total number of positions deduplicated: %i" %
               processor.UMIClusterer.positions)
        if processor.UMIClusterer.positions > 0:
            info("Mean number of unique UMIs per position: %.2f" %
                   (float(processor.UMIClusterer.total_umis_per_position) /
                    processor.UMIClusterer.positions))
            info("Max. number of unique UMIs per position: %i" %
                   processor.UMIClusterer.max_umis_per_position)
        else:
            warn("The BAM did not contain any valid "
                   "reads/read pairs for deduplication")

    Stop()


if __name__ == "__main__":
    sys.exit(main(sys.argv))
```
